# Dorsal root ganglia control nociceptive input to the central nervous system

**DOI:** 10.1101/2021.07.14.452325

**Authors:** Han Hao, Rosmaliza Ramli, Caixue Wang, Chao Liu, Shihab Shah, Pierce Mullen, Varinder Lall, Frederick Jones, Jicheng Shao, Hailin Zhang, David B. Jaffe, Nikita Gamper, Xiaona Du

**Author notes:** Correspondence to: Xiaona Du or Nikita Gamper.

## Abstract

Accumulating observations suggest that peripheral somatosensory ganglia may regulate nociceptive transmission, yet direct evidence is sparse. Here we show that the peripheral afferent nociceptive information undergoes dynamic filtering within the dorsal root ganglion (DRG) and suggest that this filtering occurs at the axonal bifurcations (t-junctions). Using synchronous *in vivo* electrophysiological recordings from the peripheral and central processes of sensory neurons (in the spinal nerve and dorsal root), ganglionic transplantation of GABAergic progenitor cells, and optogenetics we demonstrate existence of tonic and dynamic filtering of action potentials traveling through the DRG. Filtering induced by focal application of GABA or optogenetic GABA release from the DRG-transplanted GABAergic progenitor cells was specific to nociceptive fibers. Light-sheet imaging and computer modeling demonstrated that, compared to other somatosensory fiber types, nociceptors have shorter stem axons, making somatic control over t-junctional filtering more efficient. Optogenetically-induced GABA release within DRG from the transplanted GABAergic cells enhanced filtering and alleviated hypersensitivity to noxious stimulation produced by chronic inflammation and neuropathic injury *in vivo*. These findings support ‘gating’ of pain information by DRGs and suggest new therapeutic approaches for pain relief.

## INTRODUCTION

Current understanding of the somatosensory information processing largely assumes that peripheral nerves faithfully deliver peripherally-born action potentials to the spinal cord. The first synapse in the dorsal horn of the spinal cord is assumed to be the first major integration point for action potentials generated at the periphery. Such a view is represented by the Gate Control Theory of pain (Melzack and Wall, 1965) and its subsequent refinements and modifications (Braz et al., 2014; Duan et al., 2014; Mendell, 2014). While it has been proposed that information processing is more efficient the earlier it begins within the sensory pathway (Barlow, 1961; Nathan, 1976; Schmidt, 1972), the absence of true synaptic connections or interneurons within peripheral somatosensory nerves and ganglia reasonably led researchers to dismiss them as possible information processing sites. Despite this, growing evidence suggests that a degree of crosstalk between the peripheral fibers (Meyer et al., 1985a; Meyer et al., 1985b) or sensory neuron somata (Amir and Devor, 1996; Amir and Devor, 2000; Kim et al., 2016) might exist. Moreover, there is substantial experimental evidence that action potentials propagating from the peripheral nerve endings of nociceptive nerve terminals to the spinal cord can fail (or be ‘filtered’) at axonal bifurcation points (t-junctions) within the dorsal root ganglion (DRG) (Al-Basha and Prescott, 2019; Chao et al., 2020; Du et al., 2014; Du et al., 2017; Gemes et al., 2013; Kent et al., 2018; Sundt et al., 2015).

An intrinsic GABAergic signalling system within DRGs was recently proposed as a modulator of ganglionic filtering (Du et al., 2017). Yet, our understanding of how information can be modified within DRGs still remains sparse. Here we obtained direct *in vivo* evidence for ganglionic filtering mediated by the GABA signalling system and assessed if such filtering can be exploited to control pain. Using *in vivo* electrophysiological recordings from the peripheral and central processes of sensory neurons (in the spinal nerve and dorsal root), optogenetic manipulations, stem axon morphometry, and biophysical modelling we demonstrate that the DRG is a *bona fide* processing device controlling and modifying nociceptive signalling into the CNS. These findings support the existence of a ‘peripheral gate’ in somatosensory system and suggest new ways of how sensory ganglia can be targeted for pain control.

## RESULTS

### Action potentials induced by the excitation of peripheral nerve endings are filtered within the DRG

We first developed a method for *in vivo* electrophysiological recording of extracellular spiking activity from both the peripheral and central branches of the L5 spinal nerve of a rat. Spinal nerve (SN), DRG, and dorsal root (DR) were surgically exposed in anesthetized rat (Fig. 1A, B). SN and DR were then individually suspended on fine hook electrodes, while the DRG was exposed to direct drug application. This preparation allows *i)* synchronous measurement of the firing rates in the SN (before spikes enter the DRG) and DR (after spikes passed through the DRG); *ii)* sensory stimulation of the hind paw; *iii)* direct application of compounds or light to the DRG.

**Figure 1.**
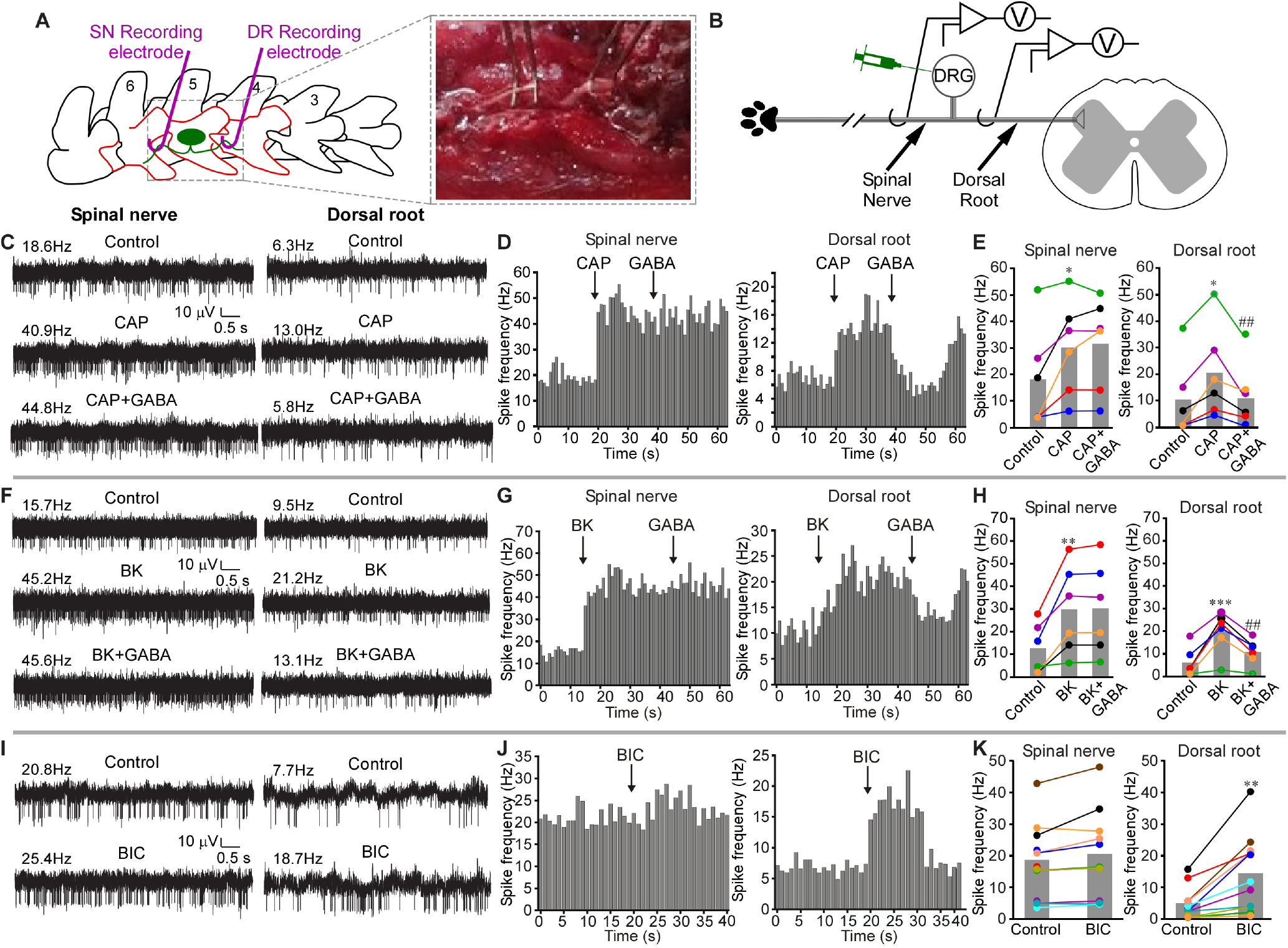
GABAergic filtering of nociceptive signals at the DRG. (**A**) Schematic of surgical exposure of the L5 spinal nerve (SN, left), L5 DRG and the dorsal root (DR, middle) in an anesthetized rat. Parts of the vertebra that are surgically removed are shown in orange. SN and DR are then individually suspended on fine hook electrodes; DRG is exposed to direct drug application. (**B**), schematic of the electrode placement. (**C**) Hindpaw injection of Capsaicin (CAP, 10 µM, 50 µl) increased firing frequency in both SN and DR branches of the nerve (middle traces, as compared to basal activity shown in the upper traces). Application of GABA (200 µM, 3 µl) to DRG reduced CAP-induced firing frequency in DR but not SN (bottom traces). (**D**) Histograms of firing frequencies from panel C. (**E**) Summary of experiments as in panel C. Two-factor repeated measures ANOVA, with factors of nerve site (SN, DR) and drug treatment (Control, CAP, CAP+GABA) revealed main effects associated with nerve site [F(1,10)=15.1; p<0.05] and drug application [F(2,9)=12.8; p<0.05], and there was significant interaction between these factors [F(2,9)=7.9; p<0.05]. Bonferroni post-hoc test: *significant difference from control (p<0.05); ^##^significant difference from CAP (p<0.01). (**F-H**) Similar to C-E, but Bradykinin (BK, 100 µM, 50 µl) was hindpaw injected, instead of CAP. (**H**) Two-factor repeated measures ANOVA: main effect associated with drug application [F(2,9)=12.0; p<0.05]; significant interaction between nerve site and drug application [F(2,9)=11.5; p<0.05]. Bonferroni post-hoc test: **,***significant difference from control (p<0.01, p<0.001); ^##^significant difference from BK (p<0.01). (**I-K**) GABA_A_ antagonist bicuculline (BIC, 200 µM, 3 µl) was applied to DRG instead of GABA; hindpaw was not stimulated. (**K**) Two-factor repeated measures ANOVA: main effects associated with nerve site [F(1,20)=7.7; p<0.05), drug application [F(1,20)=12.0; p<0.01), significant interaction between nerve site and drug application [F(1,20)=18.7; p<0.01]. Bonferroni post-hoc test: **significant difference from control (p<0.01).

In our recordings both the SN and DR usually displayed spontaneous firing activity (Fig. 1C-K), consistent with earlier reports (Janig et al., 1968; Wall, 1960). Intraplantar injection of algogenic compounds, capsaicin (a TRPV1 agonist; CAP, 10 µM, 50 µl; Fig. 1C-E) or bradykinin (BK, 100 µM, 50 µl; Fig. 1F-H), significantly increased firing frequency in both SN and DR branches of the nerve, consistent with the evoked nociceptive inputs being transmitted from the peripheral nerve towards the spinal cord. Capsaicin injection increased firing rates to 288% and 295% of basal values in SN and DR nerves, respectively; BK injection increased firing rates in SN and DR to 166% and 246%, respectively.

Recent studies suggest there is a GABAergic inhibition at the DRG, in addition to the well-accepted spinal GABAergic inhibitory network (Du et al., 2017; Obradovic et al., 2015), and that DRG neurons can themselves produce and release GABA (Du et al., 2017; Hanack et al., 2015). We thus tested how exogenous application of GABA to the DRG would affect the propagation of peripherally-induced nociceptive signals through the ganglion. Direct application of GABA (200 µM, 3 µl) to the DRG (see Materials and Methods) significantly reduced capsaicin- or BK-induced firing rates specifically in the DR, having no effect on the firing rates in the SN (Fig. 1C-H). Thus, GABAergic inhibition at the DRG can induce a prominent filtering of the throughput conduction.

Interestingly, application of the GABA_A_ receptor antagonist, bicuculline (BIC; 200 µM, 3 µl) to the DRG during continuous recording of spontaneous activity in both SN and DR (in the absence of any peripheral stimulation) significantly increased firing rate in the DR but not in the SN (Fig. 1I-K). This finding is consistent with our previous observation that application of BIC via the L5-DRG-implanted cannula *in vivo* induces nocifensive behavior towards the hind paw (Du et al., 2017). Thus, BIC is likely to attenuate tonic filtering in nociceptive fibers passing through the DRG.

The interpretation of experiments presented in Fig. 1 could be complicated by the presence of the ventral root. Even though the experiments were conducted on immobile animals under deep anesthesia, the presence of intact motor fibers in the SN allows for execution of the flexor reflex which could contribute to SN activity via the efferent and re-afferent discharge. To account for such an eventuality, we evaluated the effect of ventral root (VR) transection on the spontaneous and evoked activity in SN and DR (Suppl. Fig. 1A-J; panel J schematizes the approach). VR transection had no noticeable effect on either the SN or DR spontaneous activity (Suppl. Fig. 1A, B). Capsaicin, GABA (Suppl. Fig. 1C, D) and BIC (Suppl. Fig. 1E, F) produced effects qualitatively identical to these presented in Fig. 1: capsaicin increased firing rate in both SN and DR and GABA reduced this induced firing rate in DR but not SN. BIC increased firing rate in DR but not in SN. Interestingly, when GABA was applied to DRG on its own, without noxious stimulation of the paw, it failed to produce a significant effect on firing rate in both the SN and DR (Suppl. Fig. 1G, H). Another important observation from the experiments presented in Fig. 1 and Suppl. Fig. 1 was that firing rates in the SN were consistently higher than in the DR. Importantly, this was also true for preparations with VR transection (summarized in Suppl. Fig. 1I). Together with the fact that BIC consistently increased firing rates in the DR (with or without VR transection) these findings support the hypothesis for tonic filtering at DRG.

While VR transection eliminated the efferent input, it did not eliminate efferent fibers themselves from the spinal nerve, thus, any spurious activity in those fibers could have contributed to the SN activity and may have contributed to higher firing rates in the SN, as compared to DR. In order to eliminate efferent fibers we took advantage of the fact that VR injury causes progressive degeneration of the motoneurons and preganglionic parasympathetic neurons (PPN) (Hoang et al., 2003). Thus, we performed VR transections and allowed animals to recover; two weeks after VR transections the recordings similar to these shown in Fig. 1 and Suppl. Fig. 1 were repeated. The motor and PPN fiber degeneration was confirmed by almost complete loss of their marker, Choline acetyltransferase (ChAT) (Hoang et al., 2003) at two weeks after VR transection (Suppl. Fig. 2A). Despite the removal of efferent fibers, spontaneous firing rate in the DR was still significantly lower, as compared to SN; GABA still significantly reduced the capsaicin-induced firing rate in DR but not SN, while BIC increased firing rate in DR but not in SN (Suppl. Fig. 2B-E). Thus, under our experimental conditions, motor neuron input had no significant contribution to either spontaneous or evoked activity in SN or DR.

When we tested the effects of GABA on the capsaicin-induced firing and of BIC on spontaneous activity in SN and DR in female rats (Suppl. Fig. 3) and on male rats anesthetized with different anesthetic (isoflurane instead of pentobarbital i.p.; Suppl. Fig. 4), we obtained results qualitatively identical to these shown in Fig. 1 and Suppl. Fig. 1-2. Thus, GABA-mediated modulation of ganglionic filtering is a phenomenon reliably observed under a variety of experimental conditions in animals of either sex.

In order to better understand filtering of specific spikes in the recordings as these exemplified in Fig. 1, we developed a spike-matching method allowing to correlate SN and DR spikes (Fig. 2A; Suppl. Fig. 5-7; See Spike Sorting section of Materials and Methods for detailed description of the approach). We used this method to analyze sample datasets from recordings shown in Fig. 1. This method proved accurate (correct matching of 80 – 100 % spike pairs) in computer-generated, Poisson spike trains up to 100 Hz irrespective of the degree of filtering (not shown); the accuracy was inversely proportional to firing frequency (Suppl. Fig. 5A, B). The rate of false positive matching (FPR) was relatively low across fibre types (mostly below 7%), but faster conducting fibre types were more likely to have a higher FPR at higher spike train frequencies (Suppl. Fig. 5C).

**Figure 2.**
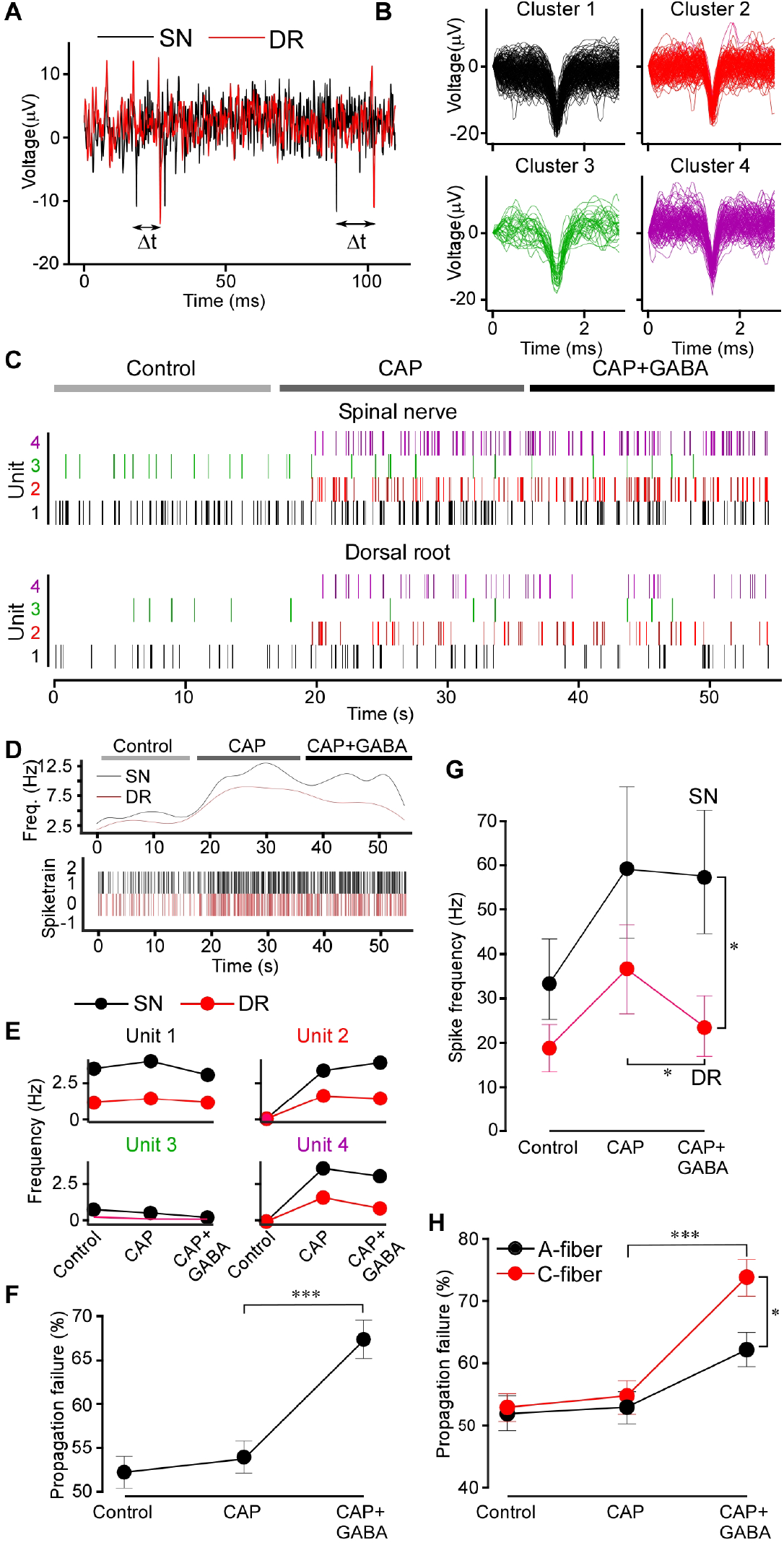
Spike analysis confirms GABAergic filtering. (**A**) Latencies between the SN and DR spikes were measured; the minimum latency defined the SN origin of a DR spike. (**B**) Extracellular spike waveforms extracted from a spinal nerve recording and clustered using WaveClus. (**C**) Raster plot for each clustered waveform (denoted as Unit) under control, CAP and CAP+GABA conditions after matching the DR spikes with these in the SN. Note: capsaicin induced firing of specific units. (**D**) Instantaneous firing frequency of the all identified units shown in C in the dorsal root and spinal nerve. (**E**) Firing rates of individual units identifies capsaicin-sensitive units. (**F**) Propagation failure (%) of all matched spike units under control, CAP and CAP+GABA conditions. One-way ANOVA: F(2,117)=17.7; p<0.001; Tukey post-hoc test: ***significant difference from CAP, p<0.001. (**G**) Average firing rates for all multi-unit nerve recordings in the dataset (n=6) in SN and DR under control, CAP and CAP+GABA conditions. Two-factor (nerve site, drug application) repeated measures ANOVA: main effects associated with drug application [F(2,10)=6.38; p<0.05], significant interaction between factors [F(2,10)=10.74; p<0.01]. Bonferroni post-hoc test: *significant difference from control (p<0.05). (**H**) Units were divided into ‘C-type’ and ‘A-type’ based on the SN-DR latency (C: <1.2 m/s; A: >1.2 m/s) and propagation failure rate analyzed as in (F). Two-factor (fiber type, drug application) mixed-effects ANOVA: main effect associated with drug application [F(2,70)=34.82, p<0.001], significant interaction between factors [F(2,72)=4.712, p<0.05]. Sidak post-hoc test: *significant difference between CAP+GABA and CAP, significant difference between A and C fibers (p<0.05).

Sorting extracellular SN spike waveforms using the WaveClus implementation of super-parametric clustering allowed us to isolate distinct spike clusters (units) and match these between SN and DR recordings. A representative experiment capturing the firing activity of four distinct units is shown in Fig. 2B-E. Another example of a similar dataset with 10 units analyzed is shown in Suppl. Fig. 6 and 7. Capsaicin-responsive units were clearly identifiable, noted by the onset of activity during application of capsaicin (Fig. 2C, E). CAP-evoked spikes were characterized by a larger amplitude on average (CAP-insensitive units: 14.5±1.5 µV; CAP-responsive units: 19.6±0.9 µV; p<0.01). However, capsaicin-responsive units could not be well distinguished by latency and spike width in this way.

Individual spike waveforms in sorted units (cluster) in the SN and corresponding average waveforms are shown in Suppl. Fig. 6B and C, correspondingly. Average waveforms of matched units in the DR are shown in the Suppl. Fig. 6D. To rule out contamination of DR units with synchronized firing of another fibre, we calculated the mean deviation (represented as a z score) of each waveform in the unit from the mean waveform of the unit (Suppl. Fig. 7A). Any spikes originating from another fibre firing in a temporally correlated way should exhibit a different waveform shape and thus be recognized as an outlier (> 3 z score). The large majority of spikes were within a z score of 3 from the unit means, suggesting our matching protocol identified homogenous and distinct DR firing units. Mean latencies for each of the 10 unit analyzed are shown in Suppl. Fig. 7B.

The mean difference in spiking units between SN and DR was significantly greater with GABA administration after CAP (Fig. 2G-H), revealing disappearance (failure) of spikes in the DR specifically upon application of GABA. We further characterized spike units as either A-fiber or C-type based on conduction velocity (<1.2 m/s; presumably C-type; >1.2 m/s, presumably A-fibers (Koltzenburg et al., 1997)). Interestingly, this revealed that C-type fibers were indeed significantly more filtered, as compared to A-type fibers, during application of GABA (Fig. 2H).

Taken together, data presented in Fig. 1, 2 and Suppl. Fig. 1-7 provide the following observations: *i)* there is a basal activity in the SN and DR even in the absence of peripheral stimulation; *ii)* the firing rates in the SN are higher than these in DR, suggesting presence of ‘tonic filtering’ of this basal activity at the DRG. *iii)* DRG-applied GABA_A_ antagonist (BIC) increased basal firing in the DR but not in the SN.

*iv)* Noxious stimulation increases firing rates in both SN and DR, but the DRG-applied GABA reduced firing in the DR specifically, thus enhancing GABAergic filtering in the nociceptive fibers. Therefore, the filtering at the DRG can also be dynamic, that is, it can be readily increased or decreased by modulating mechanisms. *v)* Exogenous GABA did not affect the basal firing rate (either in the SN or DR). This may indicate that some fibers may be already inhibited by GABA tone at the basal conditions and adding extra GABA does not affect these but inhibiting GABA channels may relieve inhibition of these ‘sensitive’ fibers. Generally, after BIC the firing rates in the DR were still somewhat lower than in the SN, hence, not all tonic filtering is necessarily GABAergic.

### Transplantation of forebrain GABAergic neuron precursors into the adult mouse DRG *in vivo* delivers an analgesic mechanism

To test how GABAergic filtering of nociceptive transmission at the DRG can be exploited *in vivo*, we adopted an approach developed by Basbaum’s group, who were able to transplant and functionally integrate into dorsal spinal cord, embryonic GABAergic progenitor cells from the medial ganglionic eminence (MGE). Transplanted MGE cells were able to compensate for the loss of spinal GABAergic inhibitory system observed in neuropathic pain models (Braz et al., 2012; Etlin et al., 2016). We transplanted embryonic MGE cells derived from VGAT-ChR2-eYFP into the L4 DRG of WT C57 mice. L4 DRG was chosen in this case as it is the major contributor to the sciatic nerve in mice (Rigaud et al., 2008). At four weeks after the DRG injection we observed numerous YFP-positive cells in the DRG sections (Fig. 3A); fluorescent cells were entirely absent in vehicle-injected control animals.

**Figure 3.**
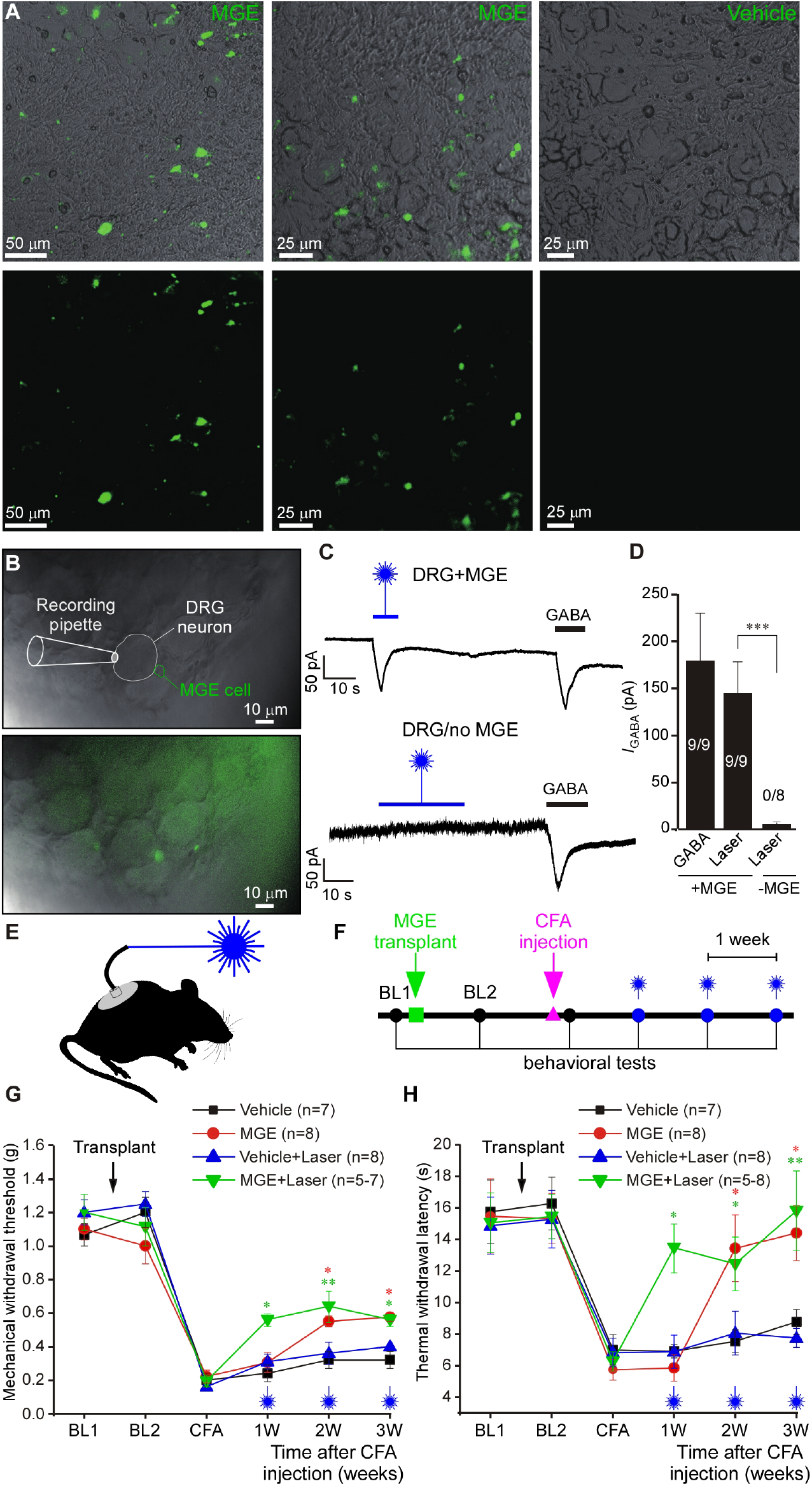
Transplantation of forebrain GABAergic neuron progenitor cells into the adult mouse DRG *in vivo* delivers an analgesic mechanism. (**A**) Fluorescence micrographs of DRG sections of mice 4 weeks after the injection with a suspension of medial embryonic eminence (MGE) cells derived from VGAT-ChR2-eYFP mice. Control mice (right images) received vehicle injections. (**B**) Bright-field and overlaid fluorescence images of ‘loosened’ DRG preparation used for patch clamp recording. The whole L4 DRG transplanted with VGAT-ChR2-eYFP MGE cells mice was extracted 4 weeks after transplantation. Recording was made from a DRG neuron juxtaposed to fluorescent MGE cell (top image) using whole-cell voltage clamp. (**C**) Top: example trace of continuous current recording (−60 mV) from the cell shown in B; stimulation with 473 nm blue laser (3 mV) induced inward current, similar in amplitude and kinetics to the current induced by perfusion of GABA (200 µM). Bottom: similar recording from a DRG of a vehicle-injected mice. (**D**) Summary for panel C; one-way ANOVA: F(2,23)=5.9; p<0.01; Bonferroni post-hoc test: ***significant difference between +MGE vs. −MGE (p<0.001). Number of recorded/responsive cells is indicated within each bar. (**E**) Schematic of the *in vivo* optogenetic DRG stimulation. (**F**) Timeline of the *in vivo* behavioral testing after the MGE cell transplantation and hindpaw injection of CFA. (**G, H**) Hypersensitivity to mechanical (G) and thermal (H) stimulation caused by hindpaw injection of CFA 2 weeks after MGE cells transplantation into L4 DRG of mice. At a time of MGE transplantation, mice were also implanted with the fiberoptic light guide. Mechanical and thermal sensitivity was measured using the von Frey and Hargreaves methods, respectively. Starting at one week after the CFA injection measurements were performed while stimulating the L4 DRG with 473nm laser. Black and blue symbols denote control mice DRG-injected with vehicle without and with optogenetic stimulation, respectively. Red and green symbols denote MGE-transplanted mice without and with optogenetic stimulation, respectively. BL1: baseline before transplantation; BL2: baseline after transplantation; CFA: 1 day after the plantar injection of CFA. **G**, Three-factor (MGE vs vehicle, time after CFA, laser stimulation) ANOVA: main effects associated with MGE transplantation [F(1,24)=50.9; p<0.001], time after CFA [F(2,23)=6.6; p<0.01]; laser stimulation [F(1,24)=8.9; p<0.01]. Bonferroni post-hoc test: red* indicate the difference between MGE group and vehicle group within the corresponding time point; green* indicate the difference between MGE with laser stimulation group and vehicle with laser stimulation group; *p<0.05, **p<0.01, ***p<0.001. **H**, Three-factor (MGE vs vehicle, time after CFA, laser stimulation) repeated measures ANOVA: main effects associated with MGE transplantation [F(1,24)=37.4; p<0.001], time after CFA [F(2,23)=6.1; p<0.01], laser stimulation [F(1,24)=2.4; p=0.12]; significant interaction between time and laser stimulation [F(2,23)=2.5; p=0.09] and between MGE and laser stimulation [F(2,23)=3.2; p=0.08]. Bonferroni post-hoc test: red* indicate the difference between MGE group and vehicle group within the corresponding time point; green* indicate the difference between MGE with laser stimulation group and vehicle with laser stimulation group; *p<0.05, **p<0.01, ***p<0.001.

In order to confirm that transplanted MGE cells can function as GABAergic neurons within DRG, we performed patch-clamp recordings from the DRG neurons juxtaposed to the MGE cells (Fig. 3B-D) using ‘loosened’ whole L4 DRGs from mice pre-injected (4 weeks) with the VGAT-ChR2-eYFP-expressing MGE (see Materials and Methods). Stimulation of the ganglion with the 473 nm blue light induced inward currents in 9/9 DRG neurons, which were in close juxtaposition with MGE cells (Fig. 3B-D). These same neurons also responded to perfusion of 200 μM GABA with very similar inward currents. In contrast, DRG neurons from vehicle-injected mice never responded to blue light (0/8 neurons) but these did respond to GABA (Fig. 3C, D). These results suggest that *i)* implanted MGE progenitor cells can survive and maturate to produce GABA-releasing neurons in DRG; *ii)* stimulus-induced release of GABA by resident neurons can induce a response in neighboring neurons.

Next, we tested if optogenetic release of GABA from the implanted MGE cells can alleviate hypersensitivity to noxious stimuli in chronic pain models. In these experiments a fiber-optic light guide was implanted into the DRG immediately after the MGE cells transplantation (Fig. 3E; Materials and Methods). Chronic inflammation with hind paw injection of complete Freund’s adjuvant (CFA, 20 µl) induced significant hypersensitivity to mechanical and thermal stimuli (Fig. 3F-H). We then performed mechanical (Fig. 3G) or thermal (Fig. 3H) sensitivity tests while stimulating ipsilateral L4 DRG with blue light. Optical stimulation significantly reduced both types of hypersensitivity in MGE-injected mice. Interestingly, starting from the second week after the CFA injection, both mechanical and thermal hypersensitivity in the MGE-implanted mice started to recover even in the absence of optogenetic stimulation and by the 3^rd^ week after the CFA injection blue light stimulation no longer produced any additional analgesic effect (Fig. 3G, H). We hypothesized that a buildup of tonic GABA release from the transplanted MGE cells in DRG could be responsible for the light-stimulation-independent recovery of the CFA-induced hypersensitivity. This hypothesis was corroborated in experiments, similar to the ones presented in Fig. 3G, H, but in which no optogenetic stimulation was used, to avoid inducing any stimulus-induced GABA release (Suppl. Fig. 8A, B). We also utilized a chronic constriction injury (CCI) model of neuropathic pain in similar experiments (Suppl. Fig. 8C, D). In both models hypersensitivity developed in control (vehicle-injected) and MGE-implanted mice. However, the latter group displayed significantly quicker and more complete recovery. Collectively, these data suggest that DRG-implanted MGE cells can be stimulated to release GABA locally *in vivo* and that such release produces analgesic effect.

### Optogenetic release or direct application of GABA to the DRG enhances filtering of spikes triggered by noxious but not innocuous stimuli

We hypothesized that the analgesic effect of MGE cells transplanted into the DRG is mediated by GABAergic filtering of pro-nociceptive spikes at the DRG. To test this we used an approach similar to that used in Fig. 1 and Suppl. Fig. 1-7, but instead of applying GABA, we stimulated L4 DRG with blue laser light (Fig. 4A). Optogenetic DRG stimulation (3 - 4 weeks after MGE transplantation) gave rise to qualitatively very similar effects to those produced by application of GABA. Firing induced by the hind paw injections of capsaicin (10 µM, 20 µl, Fig. 4B, D) or BK (100 µM, 20 µl, Fig. 4C, E) was significantly inhibited by the light stimulation in DR but not SN. The optogenetic suppression of firing in DR was evident immediately upon application of blue light (Fig. 4B, C; lower right panels).

**Figure 4.**
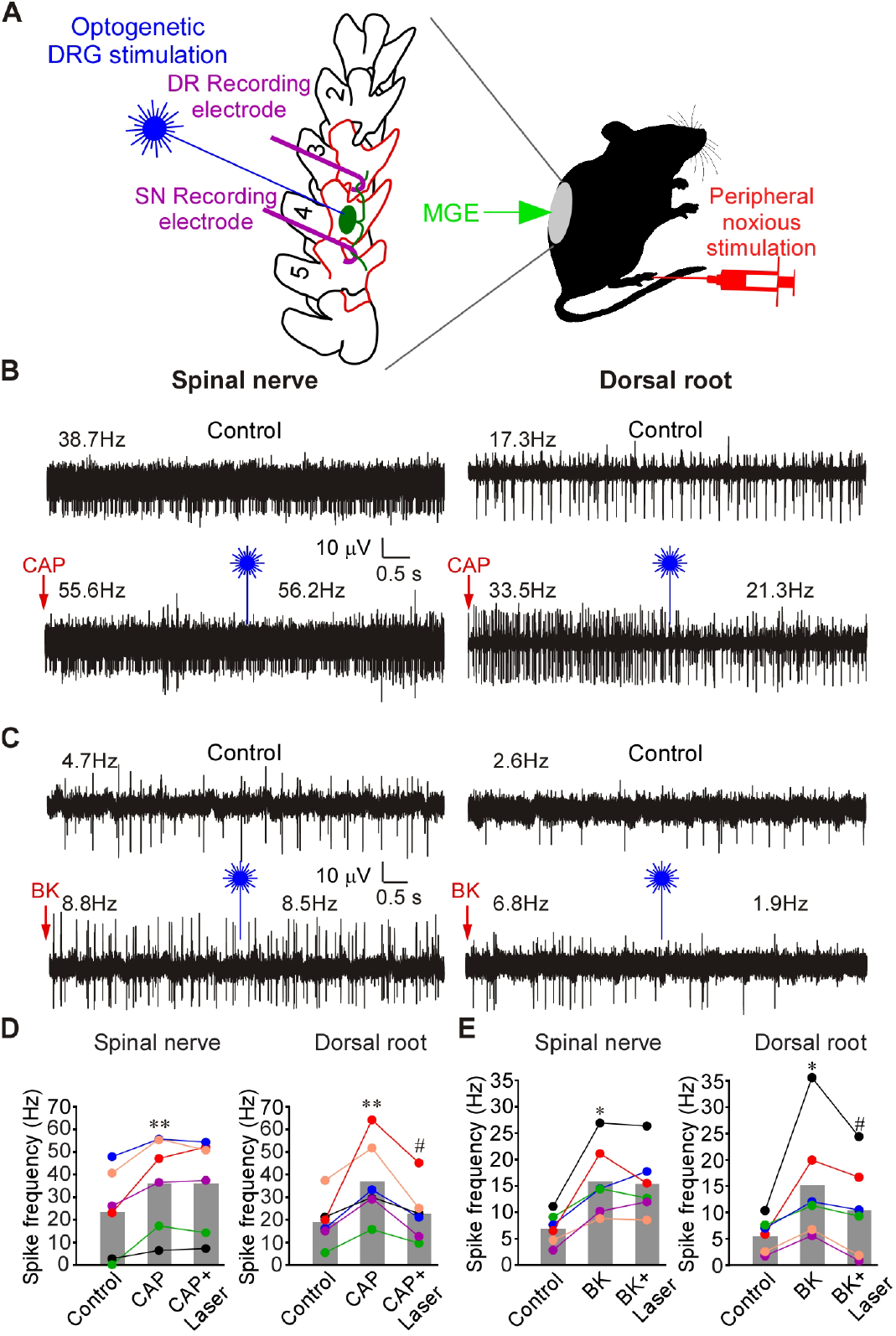
Optogenetic activation of MGE cells in DRG reduces frequency of firing induced by CAP or BK in the DR but not in the SN. (**A**) Schematic of the recording paradigm: mice were DRG-transplanted with VGAT-ChR2-eYFP MGE cells 3-4 weeks before recordings; recordings were done as shown in Fig. 1B (but from L4 DRG), supplemented with optical stimulation of the exposed DRG. (**B**) Hindpaw injection of CAP (10 µM, 20 µl; indicated by red arrow) increased firing frequency in both SN and DR. Application of 473 nm laser light to DRG (indicated by blue symbol) acutely reduced CAP-induced firing frequency in DR but not SN (bottom traces). (**C**) Experiment similar to A, but BK (100 µM, 20 µl) was hindpaw injected, instead of CAP. (**D**) Summary of panel B. Two-factor (nerve site, drug application) repeated measures ANOVA: main effect associated with drug treatment [F(2,9)=11.4; p<0.05] and significant interaction between nerve site and treatment [F(2,9)=7.6; p<0.05]. Bonferroni post-hoc test: **significant difference from control (p<0.01); ^#^significant difference from CAP (p<0.05). (**E**) Summary of panel C. Two-factor (nerve site, drug application) repeated measures ANOVA: main effect associated with nerve site [F(1,10)=6.7; p<0.05] and significant interaction between nerve site and treatment [F(2,9)=9.5; p<0.05]. Bonferroni post-hoc test: *significant difference from control (p<0.05); ^#^significant difference from BK (p<0.05).

Application of noxious heat (60° C water; Suppl. Fig. 9A, B) and noxious cold (ice; Suppl. Fig. 9C, D) induced a significant increase of firing frequencies in both SN and DR and optogenetic DRG stimulation significantly inhibited firing rates in DR but not in SN. We then tested innocuous and noxious mechanical stimulation. Air puffs and subthreshold (4g) von Frey filament stimulation (Suppl. Fig. 9E-H) increased firing in both SN and DR, presumably via the activation of low threshold mechanoreceptors in the skin (Abraira and Ginty, 2013; Li et al., 2011). Interestingly, blue light illumination of DRGs did not significantly affect the firing frequency in either SN or DR in these experiments (Suppl. Fig. 9E-H).

Noxious mechanical stimulation of the paw with a blunt glass needle also significantly increased firing frequencies in both SN and DR. In this case optogenetic stimulation substantially inhibited firing in DR, but not in SN (Suppl. Fig. 9I, J). Thus, it appears that GABAergic filtering in DRG predominantly exists in nociceptive fibers.

Next, we performed a set of experiments, similar to that shown in Suppl. Fig. 9 but in naïve rats and with the direct injection of GABA into the DRG instead of optogenetic stimulation (Suppl. Fig. 10). There was a pattern similar to that observed with optogenetic stimulation of MGE cells in mice. Firing induced by noxious heat (Suppl. Fig. 10A, B) and noxious cold (Suppl. Fig. 10C, D) was selectively inhibited in the DR but not in the SN by the DRG-applied GABA. No significant effects of DRG-applied GABA were seen when firing was induced by innocuous air puffs or subthreshold von Frey hairs (Suppl. Fig. 10E-H). In contrast, firing induced by the noxious needle prick was reduced in the DR but not in the SN (Suppl. Fig. 10I, J). Striking similarity of the effects of exogenous GABA and optogenetic GABA release in DRG strongly suggest that *i)* the GABAergic system modulates filtering efficacy of the ganglia; *ii)* the filtering is most efficient in nociceptive fibers and *iii)* such filtering is the most plausible explanation of the analgesic effect of MGE cells transplanted to the DRG; *iv)* GABAergic filtering exists in both rats and mice.

To test this further we performed single-unit recordings (Fig. 5A-G, schematized in panel G). Solutions (200 μM GABA, 1 μM TTX or saline; all in 3 μl volume) were applied to the DRG by micropiettor and firing was induced before or after the injection by a stimulus train delivered by a stimulating electrode placed in SN while recordings were made from the mechanically isolated teased DR bundles. A-type and C-type spikes were distinguished by the conduction velocity (A fibers >1.2 m/s; C fibers <1.2 m/s (Koltzenburg et al., 1997)). The firing in both fiber types was blocked by 1 μM TTX applied to the DRG (Suppl. Fig. 11A, B, E). In single-unit recordings evoked stimuli propagated reliably in both fiber types under control conditions with only occasional failures (basal failure rate in A fibers: 5.6±2.4%, n=8; C fibers: 15.8±2.4%, n=9; Fig. 5A-F), which is consistent with earlier report (Al-Basha and Prescott, 2019). At 10 Hz stimulation DRG-applied GABA significantly increased failure rate in C fibers (to 65.0±3.1%, p<0.001) but had no significant effect in A fibers (Fig. 5A, C, E). Even when stimulation frequency was increased to 50 and 100 Hz, DRG applied GABA still failed to affect spike propagation in A fibers (Suppl. Fig. 11C, D, F). DRG injection of vehicle (saline) had no effect on filtering (Fig. 5B, D, F). Due to technical difficulties of these recordings not all of these were long enough to reliably analyze failure rate but amongst all the recordings made, 90% (38/42) of C fibers and only 17% (11/63) of A-type fibers displayed GABA-induced t-junctional spike failure, while in 100% of both fiber types failure was induced by TTX (Suppl. Fig.10E). Taken together, single-unit recordings revealed that DRG application of GABA selectively increases filtering of C-fiber activity.

**Figure 5.**
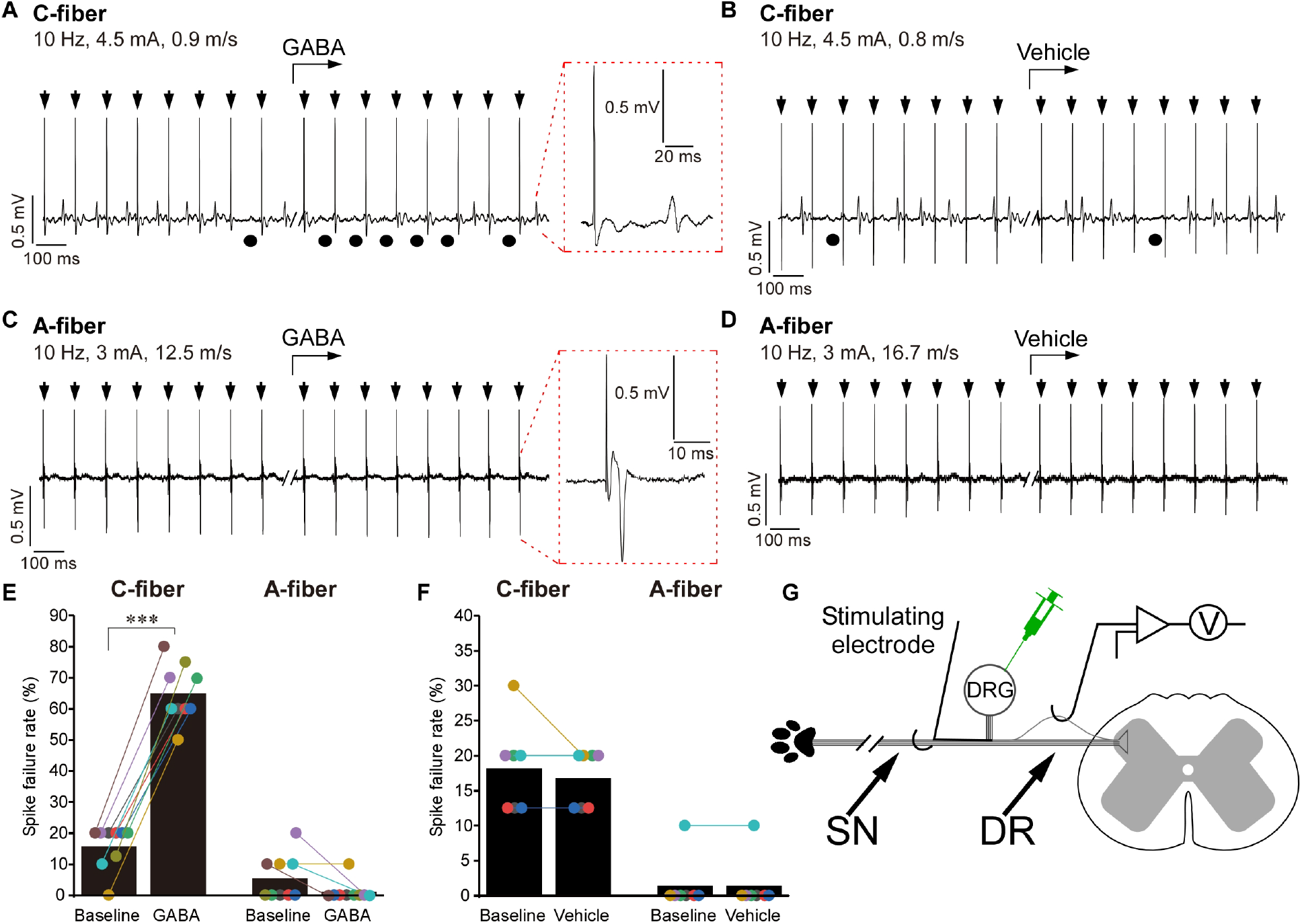
Single-unit recordings: DRG application of GABA induces spike failure at the DRG, which is specific to C fibers. (**B-D**) Example traces of *in vivo* single unit recording from the DR process of a rat C fiber (A, B) or A fiber (C, D); stimulus electrode is placed in the spinal nerve (schematic of the experiment is shown in panel D). Parameters of stimulation and conduction velocity are indicated above the trace. GABA (200 µM, 3 µl; A, C) or vehicle (3 μl saline; B, D) were applied to the DRG by a micropipettor at the time point indicated by the bent arrows. Stimulus artifacts are indicated by black arrows. Black circles under the traces indicate failed spikes. Spike waveform is shown on the extended time scale within the dotted red boxes. (**E**) Spike failure rate before and during application of GABA is analyzed for C and A fibers; n=9 (one recording per animal) for both fiber types; Kruskal-Wallis ANOVA: H(3)=28.2, p<0.001; Mann-Whitney post-hoc test: ***significant difference from control (p<0.001). (**F**) Similar to panel E but spike failure rates before and after the application of vehicle control are analyzed; n=7 for both fiber types. No significant effects of the vehicle were found (Kruskal-Wallis ANOVA). (**G**) Schematic of the single unit recording paradigm.

### Does the soma have more influence over the t-junction in C-type as compared with A-type fibers?

DRG neurons are pseudo-unipolar and axonal bifurcation (t-junction) is potentially a major site of spike filtering in DRG due to impedance mismatch (Debanne et al., 2011; Du et al., 2017; Gemes et al., 2013; Goldstein and Rall, 1974; Kent et al., 2018; Luscher et al., 1994b; Sundt et al., 2015). Why is GABA-induced filtering more efficient in the C-fibers, as compared to A-fibers? One possibility is the different length of the stem axon (from the soma to the t-junction) and, thus, the electrotonic influence of the soma on the t-junction and spike propagation: the longer the stem axon, the poorer the coupling (Du et al., 2014; Du et al., 2017; Sundt et al., 2015). To our knowledge, there has been no systematic analysis of stem axon lengths in mammalian DRG neurons, although drawings by Ramon y Cajal depict much shorter stems in small-diameter neurons, as compared to larger ones (Ramón y Cajal, 1909). In addition, larger neurons are often depicted having stems with a winding ‘glomerular’ section, extending the length (Matsuda et al., 2005; Ramón y Cajal, 1909; Ranson, 1912). In order to assess stem length in C-type *vs*. A-type fibers, we cleared (Renier et al., 2016) rat whole DRG mounts and immunostained them with the C-fiber marker, peripherin, and A-fiber marker, neurofilament-200 (NF-200). We then performed light-sheet microscopy of entire ganglia (Movie S1) and measured the stem axon lengths of peripherin-positive and NF-200 positive neurons (Movie S2; Fig. 6A-B). Consistently, peripherin labelled neurons with much smaller somatic diameter, as compared to NF-200 positive neurons (26.1±0.4 μm *vs*. 42.4±0.8 μm, p<0.001; Fig. 6A, E; Suppl. Fig.12). Stem axon diameters of peripherin-labelled neurons were also consistently smaller (1.34±0.03 μm *vs*. 2.1±0.1 μm, p<0.001; Fig. 6D; Suppl. Fig. 12). Of note, axonal diameters for NF-200 positive neurons reported here do not include myelin and are in good agreement with previous literature (Suaid et al., 2016). Peripherin labelled neurons displayed much shorter stems, as compared to NF-200 positive neurons (60.7±4.2 μm *vs*. 232.5±22.9 μm, p<0.001; Fig. 6C; Suppl. Fig. 12). While for all peripherin-positive neurons analyzed in Fig. 6C and Suppl. Fig. 12, the t-junction was reliably identified (Movie S2), it was often impossible to confidently locate the t-junctions of NF-200 positive neurons as these were too far away from the cell body. In these instances stem length was recorded as the longest traceable distance and, hence, it is an underestimation of the real stem length.

**Figure 6.**
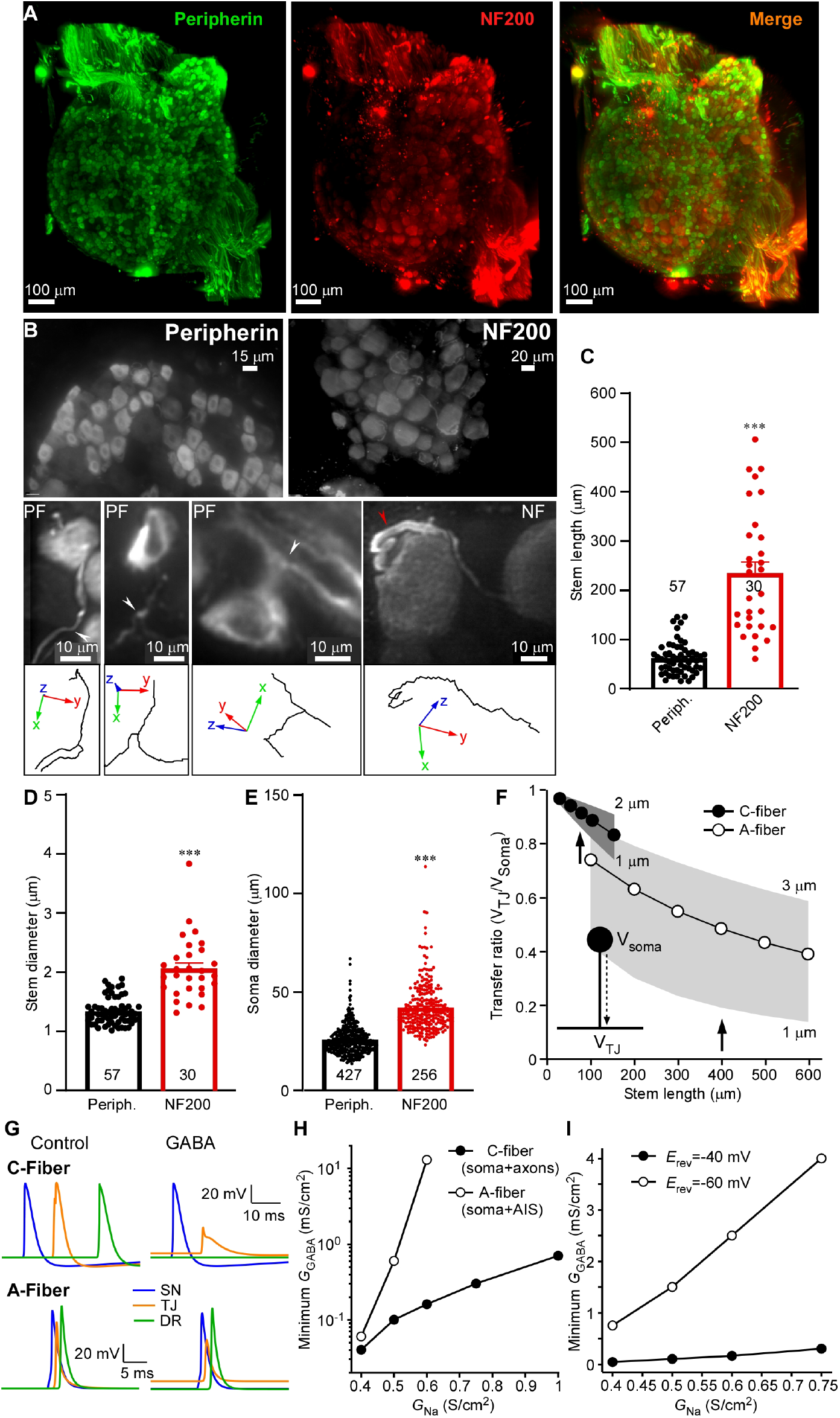
Stem axon morphology defines the efficiency of filtering in DRG. (**A-B**) Light-sheet microscopy of cleared rat DRG (see also Movie S1). C-fibers are labelled with peripherin and A-fibers are labelled with NF-200. Stem axons were traced until bifurcation point using simple neurite tracer in FIJI (Movie S2). White arrowheads indicate t-junctions, red arrowhead point at glomerular region of A-fiber stem axon which increases the length. (**C-E**) Quantification of the stem axon lengths (C), stem diameter (D) and somatic diameter (E) of the peripherin- and NF-200-labelled fibers is shown. Data are from 10 lumbar DRGs (peripherin) and 7 lumbar DRGs (NF200) from three rats; n number is given within each bar. Unpaired t-test: t(85)=9.8, p<0.001 (C); t(85)=9.8, p<0.001 (D); t(681)=21.6, p<0.001 (E). Per-animal analysis of data in C-F are shown in Suppl. Fig. 12. (**F-I**) Biophysical modeling of sensory neurons. (**F**) Voltage attenuation from the soma to the t-junction (TJ) measured as the transfer ratio (V_TJ_/V_Soma_) for both C- and A-fiber models, across a physiological range of stem axon lengths and diameters. Note that there is overlap only at the shortest range of A-fiber model lengths and widest diameters. Thus, voltage attenuation is predicted to be greater for A-fiber than C-fiber stem axons. (**G**) Top traces are from a C-fiber model that exhibits spike propagation in the absence of GABA_A_ receptor activation and GABA-sensitive spike failure. In this case, G_Na_ across the model was lowered (0.4 S/cm^2^) and GABA receptors (0.04 mS/cm^2^) expressed in the soma and axons. In contrast, traces from an A-fiber model configured to be insensitive to the effects of GABA; higher G_Na_ (0.75 S/cm^2^) and GABA expression (0.02 mS/cm^2^) restricted to the soma and AIS. (**H**) Across a wide range of G_Na_ densities the C-fiber model was more GABA-sensitive to spike failure compared with the A-fiber model. In contrast, the A-fiber model with GABA receptors restricted to the soma and AIS is less sensitive. Except for the lowest density of G_Na_, a higher density of GABA receptor activation is required to induce spike failure. Moreover, at higher G_Na_ densities (greater than 0.6 S/cm^2^ in the model) GABA receptor activation at any level no longer prevents spike propagation. (**I**) In a C-fiber model where E_Cl_ was set to the resting potential, and didn’t depolarize membrane potential, the minimum GABA conductance required to block spikes was greater than when E_Cl_ = −40 mV across a range of G_Na_, consistent with the requirement for inactivation of Na^+^ channels to lower the safety factor for spike propagation.

We hypothesized that one reason for why GABAergic control of DRG filtering is less effective in A-fibers compared to C-fibers may be due to the differences in stem axon length and the influence of somatic conductance load. Our previous computational modeling suggested that in a C-fiber with 75 μm stem axon, activation of somatic GABA_A_ channels can indeed cause a failure of action potential to propagate through the t-junction due to a combination of the impedance drop and sodium channel inactivation (Du et al., 2017). But even though data presented in Fig. 6A-C indicate that the stem axon of C-fibers is, on average, at least 3 times shorter than that of A-fibers, myelination could possibly compensate for the longer distance. Additionally, the larger diameter of A-fiber neurons, and greater conductance load, might also compensate for a longer stem axon. Using a computational model of an A-fiber neuron, we examined the relationship between somatic conductance and stem axon length. We constructed a minimal model of an A-type neuron (see Materials and Methods) containing a limited repertoire of voltage-gated channels and a geometry consistent with the parameters obtained from the light-sheet morphometry of the NF-200 positive neurons and compared it to a model with a C-fiber morphology (Du et al., 2017).

We first examined how potential at the soma influences the t-junction between the two models. We calculated the voltage transfer ratio, the fractional potential reaching the t-junction produced by a DC potential generated at the soma, for a passive model as a function of stem axon length and diameter for the two models (Fig. 6F). In C-fibers, the transfer ratio was greater than 0.75 for a physiological range of stem lengths, while it was less than 0.6 for stem axon lengths measured from A-type neurons. For A-fibers, electrotonic control was comparable to the C-fiber model when stem axon lengths were limited to the shorter end of the range and stem axon diameter was wider. Thus, based simply on electrotonic distance, modeling predicts that the soma of a C-fiber neuron should have more influence on the t-junction than for an A-type neuron soma.

We then examined the propagation of spikes initiated in the most distal portion of the SN axon through the t-junction into the DR axon. For a myelinated A-fiber model spikes at the nodes of Ranvier in SN and DR axons were 86 and 91 mV in amplitude, respectively (nodal G_Na_ = 0.2 S/cm^2^). Conduction velocity in the SN axon was 7 m/s for an internode distance of 150 μm. In our unmyelinated C-fiber model spikes were 89 mV in amplitude in SN and DR axons (G_Na_ = 0.4 S/cm^2^), and SN axon conduction velocity was 0.2 m/s (Fig. 6G, left). In both models, the repertoire of active conductances was limited to only TTX-sensitive Na^+^ (Herzog et al., 2001) and delayed rectifier K^+^ (Sheets et al., 2007) channels, allowing us to compare the effects of geometry between the two models. For the A-fiber model we started with a stem axon of 400 μm and 2 μm diameter, and the C-fiber model started with a stem axon of 75 μm and diameter 1.35 μm. Conduction through the t-junction across a wide range of physiologically-relevant parameter space was not influenced by firing at the soma, consistent with earlier models (Amir and Devor, 2003).

To assess the potential effect of GABA_A_ receptor activation on spike propagation through the DRG, a chloride conductance (G_Cl_) was introduced with E_Cl_ = −40 mV (Liu et al., 2010). When GABA_A_ receptors were introduced only to the soma, an increase in somatic G_Cl_ depolarized the t-junction and, in turn, reduced local spike amplitude in both models. GABA_A_ receptor activation at the soma only could block spike propagation in both models, but the minimum conductance required to block propagation for the A-fiber model was 0.25 mS/cm^2^ (25 nS net conductance accounting for area) compared with 0.5 mS/cm^2^ (5 nS net conductance) for the C-fiber model (G_Na_ = 0.5 S/cm^2^). Currently, there is no evidence that GABA_A_ receptors are limited to the soma and it is possible that GABA_A_ receptors are expressed on both the soma and axons of sensory neurons, although the total conductance provided is likely to be less at nodes of Ranvier. The greatest sensitivity to GABA_A_ receptor activation for the C-fiber model was best achieved when GABA_A_ receptors were expressed throughout the model (Fig. 6G, right). In this case, the minimum G_Cl_ needed to block spikes (i.e. threshold) was substantially lower in the C-fiber model, over a wide range of Na^+^ channel densities (Fig. 6H). As expected, for A-fiber models when the stem axon length was systematically varied, we found that longer stem axons had less influence on t-junction potential, and in turn Na channel inactivation (Supp. Fig. 13).

We expect an increase in G_Cl_ to not only depolarize the t-junction, but also to act as a shunt via the increased membrane conductance, which would be expected to affect local impedance. In order to separate the effects of shunting versus depolarization, in the C-fiber model we compared the minimum G_Cl_ needed to block spikes at two equilibrium potentials for chloride (E_Cl_); first the model was set to realistic E_Cl_ = −40 mV and then it was changed the value of the resting potential (−60 mV). As shown in Fig. 6I, with only shunting (E_Cl_ = −60 mV), greater G_Cl_ was required to block spikes, and was proportional to G_Na_. This indicates that membrane depolarization, and the subsequent voltage-dependent inactivation of Na^+^ channels, contributes to the lower safety factor for spike propagation in the presence of GABA_A_ receptor activation. Indeed, when we examined the inactivation state variable for the Na^+^ conductance, the fractional available Na^+^ conductance at the t-junction decreased with increasing somatic G_Cl_ (Suppl. Fig. 13). Taken together, the morphometry of DRG neuron stem axons and our biophysical models support the hypothesis that the longer stem axons of A-fibers contribute to limiting the influence of somatic GABA_A_ conductance on spike filtering. That said, expression of G_Cl_ in the axons (stem, SN, and DR) of the models was more effective at inactivating Na^+^ channels, compared to when they were only expressed at the soma (Suppl. Fig. 13, bottom graphs in panels A and B). Additionally, greater shunting at, and proximal to, the t-junction lower the safety factor for spike propagation. Thus, under conditions with homogenous distribution of GABA_A_ channels in the soma and axons, and the uniform concentrations of GABA in the extracellular space, the axonal GABA_A_ channels would have stronger influence over the t-junctional filtering than somatic GABA_A_ channels. However, if GABA is released onto the soma mostly (i.e. across the satellite glia septum or in an autocrine way), then the effectiveness of electrotonic coupling of the soma to the t-junction would again be dependent on the stem axon length. In this vein, long stem axons of A fibers may physically remove the t-junctions away from the somatic GABA release area, making axonal GABA_A_ channels near the t-junction less relevant.

### Evidence for tonic release of GABA within the DRG

Our paired recordings from SN and DR suggest the existence of tonic filtering at the DRG. At least a proportion of this tonic filtering is GABAergic (presumably due to a GABA tone), since BIC increases firing rate in the DR (Fig. 1G, H; Suppl. Fig. 1-4), moreover, when injected to the DRG *in vivo*, BIC induces pain-like behavior (Du et al., 2017)). In non-nociceptive fibers this tonic filtering is much less pronounced (Fig. 5). Here we focused on the GABAergic mechanism; our previous data suggested that some DRG neuron cell bodies are capable of releasing GABA upon stimulation (Du et al., 2017). Another recent study reported robust activity-dependent somatic vesicle release from DRG neurons (Chai et al., 2017). To investigate mechanisms of GABA release by DRG neurons we developed a method for measuring exocytosis of GABA-containing vesicles based on the live uptake of luminal (C-terminal) antibody against vesicular GABA transporter VGAT (Fig. 7A). N-terminus of VGAT, inserted in the neurotransmitter vesicle membrane, faces the cytoplasm while the C terminus resides in the vesicle lumen (Martens et al., 2008). During exocytosis and subsequent recycling of a vesicle, luminal VGAT epitopes are temporarily exposed to the extracellular milieu. During such an exposure, antibodies that recognize these epitopes can bind to these and become trapped and subsequently internalized by endocytosis (Fig. 7A; (Martens et al., 2008)). Antibodies against the N-terminus of VGAT should not be entrapped in this way as N-terminus of VGAT remains cytosolic at all times. Depolarization of cultured DRG neurons with extracellular solution containing 100 mM KCl induced robust uptake of C-terminal (luminal) but not N-terminal (cytosolic) VGAT antibody by DRG neurons (Fig. 7B, C; quantified as proportions of neurons stained with the C-terminal VGAT antibody). The depolarization-induced C-terminal VGAT antibody uptake was significantly reduced (but not abolished – in good agreement with (Chai et al., 2017)) by the removal of extracellular Ca^2+^ (Fig. 7B, C). Interestingly, even in the absence of depolarization, there was a significant number of neurons that took up C-terminal VGAT antibody (Fig. 7B top panel; C), hinting at spontaneous exocytosis of VGAT-containing vesicles. The N-terminal VGAT antibody was not taken up by DRG neurons (Fig. 7B, bottom panel; C), even though this same antibody labeled permeabelized DRG neurons well (Du et al., 2017).

**Figure 7.**
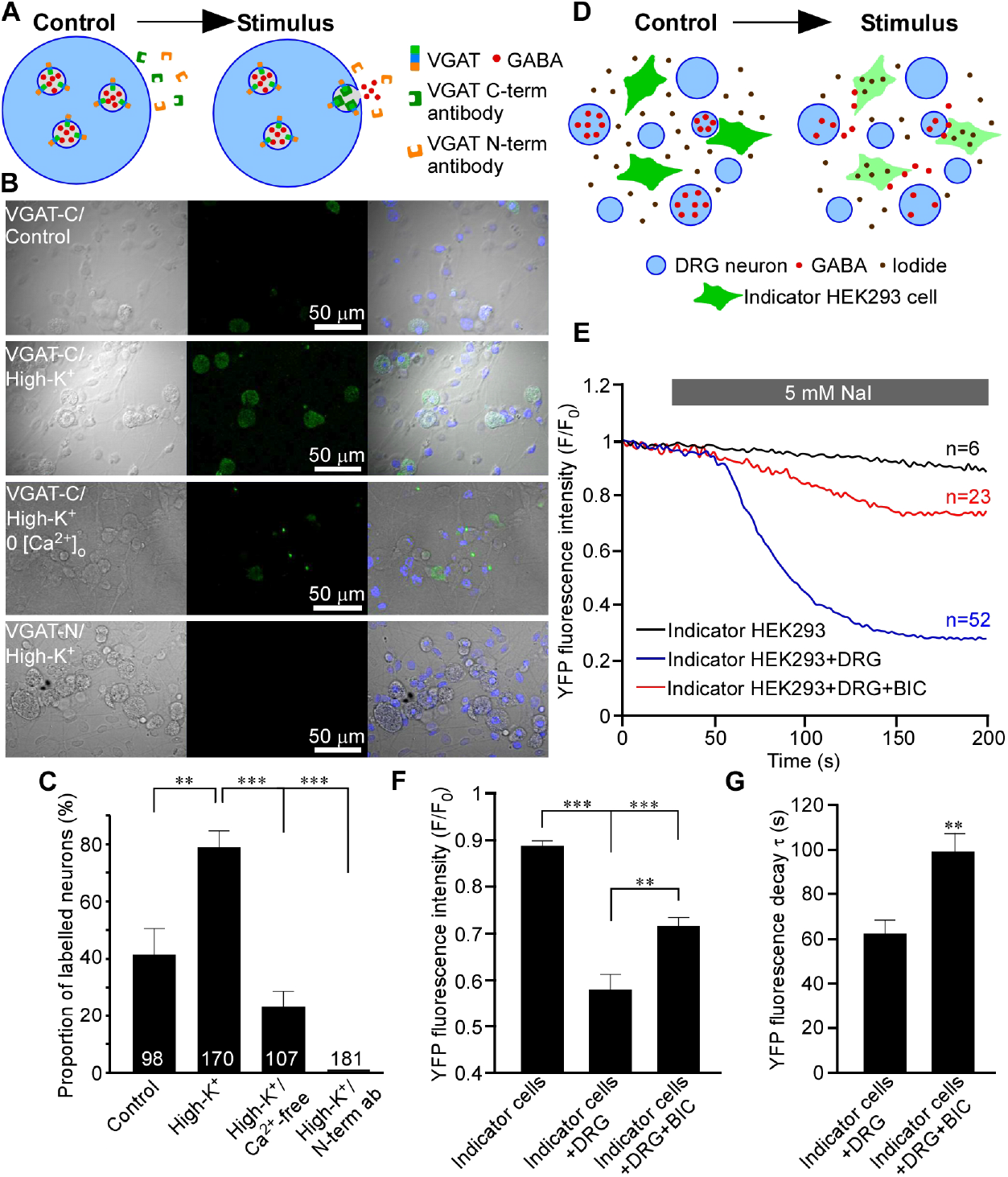
Activity-dependent and spontaneous GABA release of GABA from DRG neurons. (**A-C**) Vesicular GABA release from DRG neurons studied with luminal VGAT antibody uptake. (**A**) Schematic of activity-dependent uptake of luminal (C-terminal) VGAT antibody. (**B**) Bright-field (left), fluorescence (middle) and overlaid images of DRG culture incubated in the presence of either a luminal VGAT antibody (C-term; three upper rows) or a cytoplasmic (N-term; bottom row) antibody. Cells were incubated in the control extracellular solution (upper row) or depolarized by extracellular solution containing 100 mM KCl (second from the top and the bottom rows). In the second from the bottom row Ca^2+^ was excluded from the extracellular solution. (**C**) Summary of experiments like these shown in B (N=3-5, number of neurons is indicated within the columns); quantified is the percentage of stained neurons **p<0.01, ***p<0.01 (Fisher’s exact test with Bonferroni correction). (**D-G**) GABA release from DRG investigated with the use of indicator HEK293 cells. (**D**) Schematic of the experiment: HEK293 cells transfected with α1, β2 and γ2 subunits of GABA_A_ receptors and a halide-sensitive EYFP mutant (H148Q/I152L; EYFP-QL) are co-cultured with DRG neurons and imaged in the presence of 5 mM extracellular iodide. When GABA is present in the extracellular solution, GABA_A_ channel opening causes I^-^ influx and EYFP-QL fluorescence quenching. (**E**) Application of NaI induced bicuculline-sensitive EYFP-QL quenching only when indicator HEK293 cells are co-cultured with DRG but not in monoculture; representative traces are shown for each experimental condition. (**F**) Summary for panel E; normalized fluorescence intensity is quantified at 200 s of each recording. Kruskal-Wallis ANOVA: H(2)=33.6, p<0.001; Mann-Whitney post-hoc test: significant difference between indicated groups, **p<0.01, ***p<0.001 (**G**) Quantification of the kinetics of the EYFP-QL fluorescence quenching in experiments shown in E; individual recordings were fit with exponential function and time constants (τ) of the fluorescence decay were analyzed. Mann-Whitney test: U=786, p<0.01. In E-F, data from three independent experiments for each condition are shown, number of cells analyzed is indicated above each trace in panel E.

Additionally, we used live imaging to optically monitor GABA release from the DRG neurons in culture. We transfected HEK293 cells with α_1_, β_2_, and γ_2_ GABA_A_ subunits and a halide-sensitive EYFP mutant (H148Q/I152L; EYFP-QL). The fluorescence of EYFP-QL is quenched by iodide and since Cl^-^ channels (e.g. GABA_A_) are permeable to this anion, EYFP-QL fluorescence quenching by I^-^ can be used to monitor Cl^-^ channel activation (Galietta et al., 2001; Johansson et al., 2013; Namkung et al., 2011; Shah et al., 2020). We co-cultured these ‘GABA indicator’ HEK293 cells with DRG neurons and measured EYFP-QL fluorescence quenching induced by the addition of 5 mM NaI to the extracellular solution (Fig. 7D). NaI induced robust EYFP-QL fluorescence quenching only when the indicator cells were co-cultured with DRG, not in the monoculture (Fig. 7E). Moreover, this quenching in the DRG/indicator cell co-culture was significantly reduced in the presence of 50 μM BIC (Fig. 7E, F). BIC also significantly slowed down the kinetics of the EYFP-QL fluorescence intensity decay (Fig. 7G). These data further support existence of GABA tone within the sensory ganglia.

## DISCUSSION

There is substantial experimental evidence indicating that DRGs regulate the flow of sensory information to the CNS (Al-Basha and Prescott, 2019; Chao et al., 2020; Du et al., 2014; Dun, 1955; Gemes et al., 2013; Luscher et al., 1994a; Stoney, 1985; Tagini and Camino, 1973). Consistent with this, electrical stimulation of DRGs provides strong analgesia even to the most intractable types of chronic pain in humans (Deer et al., 2013; Hunter et al., 2019; Liem et al., 2013). Simulations and experiments point to axonal t-junctions within DRGs as biophysically amenable sites for peripheral filtering/gating (Du et al., 2014; Du et al., 2017; Kent et al., 2018; Luscher et al., 1994a; Sundt et al., 2015).

Action potential processing within DRGs can be modulated by GABAergic signalling. Thus, DRG neurons can produce and release GABA (Du et al., 2017; Hanack et al., 2015) and GABA delivered directly into the DRG has prominent analgesic action (Du et al., 2017; Obradovic et al., 2015). Our main findings here are threefold. First, we demonstrate that the DRG exerts a dynamic control over the throughput conduction via GABA receptor activation. Indeed, focal application of GABA, or optogenetic stimulation of GABA release from the DRG-transplanted MGE cells, specifically reduced stimulus-induced firing frequencies in the DR but not the SN in various experimental settings. Second, we show that this GABAergic filtering is much more efficient in the C-fibers, as compared to A-fibers. Third, we demonstrate that an intrinsic GABAergic inhibitory system can be engaged to scale the filtering of spikes passing through the DRG up or down, modulating nociceptive input to the CNS. Importantly, we further show that this mechanism can significantly reduce sensitivity to noxious stimuli and provide effective relief of hypersensitivity observed in chronic pain models.

The t-junctions, where the stem axon bifurcates into SN and DR axons, are expected to have a lowered safety factor for AP propagation due to the impedance mismatch (Debanne, 2004; Du et al., 2014; Du et al., 2017; Gemes et al., 2013; Luscher et al., 1994b; Stoney, 1985). Our earlier biophysical model of a C-fiber predicted that opening of somatic/peri-somatic GABA_A_ channels increases the likelihood for the failure of action potential propagation through the t-junction (Du et al., 2017). Here we provide strong experimental support to these predictions. We show that GABA receptor activation within the DRG reduces the firing rate recorded at a point after the t-junction, and has no effect on spikes entering the DRG. The most straightforward interpretation of these observations is that evoked spikes in peripheral axons fail to propagate through the DRG into DR axons when GABA system is activated in the DRG. According to earlier modeling, the failure maybe due to a combination of t-junction geometry (i.e. input impedance drop), GABA conductance shunting, and GABA-induced depolarization and subsequent voltage-gated sodium channel inactivation (Du et al., 2017). It is important to point out that in this study we focused specifically on GABAergic filtering but this does not imply that t-junctional filtering cannot be mediated or regulated by other mechanisms. For instance, activation of somatic, or stem axon/t-junction K^+^ channels has also been suggested as a mechanism promoting t-junctional spike failure (Chao et al., 2020; Du et al., 2014).

Another striking finding is that GABAergic DRG filtering is much more robust in C-, as compared to the A-fibers. As a consequence, optogenetic GABA release (Suppl. Fig. 9) or DRG injection of GABA (Suppl. Fig. 10) specifically inhibited firing produced by noxious but not innocuous stimuli. We propose that the induced filtering is more efficient in the C-fibers in part, because these have a shorter stem axon. Electrotonic coupling of the soma to the t-junction depends on the stem axon length (Du et al., 2017; Sundt et al., 2015). Drawings by Ramon y Cajal indeed suggested much shorter stems in small-diameter DRG neurons, as compared to the larger ones (Ramón y Cajal, 1909). Our light-sheet imaging of cleared DRG with labelled C- and A-type fibers established that stem axon is indeed over 3 times shorter in C fibers (Fig. 6). Our new biophysical model of an A-fiber suggests that when a stem axon is longer than 200-300 μm, opening of somatic GABA_A_ channels has less effect on the action potentials propagating through their t-junctions. Thus, this investigation may have revealed a simple but elegant principle of differential filtering of spikes that depends on the modality-specific fiber morphology.

It should also be emphasized that reduced excitability at the t-junction, not the impedance mismatch alone, contributes to GABA-dependent spike failure. Indeed, there is evidence to suggest that the density of Na channels is not uniform, at least between the soma and axons (Akopian et al., 1999; Klein et al., 2017). It has to be acknowledged that a minimal model is presented here and that other mechanisms, e.g. differential repertoires and densities of ion channels (including but not limited to GABA_A_) at the t-junctions or stem axon of different fiber types could also contribute to variations in filtering. Thus, in the absence of internodes, C fibers may have a higher GABA_A_ conductance density along their axon. There well may be other dynamic and activity-dependent differences in gating of action potentials at the t-junctions of different fiber types (Petersson et al., 2014; Tigerholm et al., 2014).

Interestingly, a recent study used single unit recordings from the DR to demonstrate that electric field stimulation of the DRG in rats, which mimicked DRG field stimulation (neuromodulation) used for analgesia in humans (Deer et al., 2013; Hunter et al., 2019; Liem et al., 2013), increased t-junctional filtering mostly in C-fibers, with little effect in A-type fibers (Chao et al., 2020). Our results are in good agreement with these findings; together these data could explain otherwise ‘paradoxical’ (but effective) analgesic approach via electrical DRG stimulation.

Two consistent, but yet to be fully understood, observations reported here require further consideration. First, there was considerable basal activity under control (no peripheral stimulation) conditions. This may arise from the activity of low-threshold, non-nociceptive neurons or from the preparation-specific injury. However, our previous observation that DRG-injected GABA antagonists in freely behaving animals induced pain-like paw flinching in the absence of noxious input (Du et al., 2017) may suggest that there is indeed some spontaneous activity in the peripheral nerve which is failing to reach spinal cord. Second, basal firing rates in the DR were almost invariantly about 50% lower, as compared to these in SN, even under conditions where efferent fibers were either cut (Suppl. Fig. 1) or physically destroyed (Suppl. Fig. 2). Single-unit recordings (Fig. 5) revealed that evoked spikes in C and A fibers do fail occasionally under basal conditions (at a rate of ~16% in C and ~6% in A fibers), which was consistent with a recent report (Al-Basha and Prescott, 2019). Thus, basal filtering does exist but when measuring evoked single-unit spikes it is much lower than ~50% mismatch seen in our multi-unit SN/DR recordings. While GABA_A_ antagonist BIC significantly reduced this mismatch indicating that at least part of it is due to tonic GABAergic inhibition, we cannot exclude that other factors may contribute to the mismatch, including e.g. some back-propagation in the SN. It is important to note however that regardless of the nature of this mismatch in basal firing, the sensory stimulation of the paw (noxious or innocuous) is seen in our SN/DR recordings as a consistent increase in the firing rates in both compartments (SN and DR). Moreover, we were able to identify individual capsaicin-induced units using our spike matching algorithm (Fig. 2, Suppl. Fig. 5-7). Thus, the activity induced by noxious stimulation is reliably detectable on the background of the basal activity. Importantly, it is this induced noxious activity, which is specifically filtered at the DRG by the GABAergic system.

Another important question - is the intrinsic GABAergic system in the DRG sufficient to impose filtering *in vivo*? Strong expression of functional GABA_A_ receptors in DRG is well recognized (reviewed in (Wilke et al., 2020)) but the ability of DRG to produce and release GABA is less well documented (although see (Du et al., 2017; Hanack et al., 2015). Transcript levels of enzymes (GAD65, GAD67) and transporters (VGAT) needed to synthesize and package GABA into vesicles are low according to transcriptomic studies (Jager et al., 2020; Usoskin et al., 2015). Yet functional proteins are detectable (Du et al., 2017; Hanack et al., 2015; Tzeng et al., 2021), as is tonic and induced GABA release (Fig. 7 and (Du et al., 2017)). It has to be pointed that a non-vesicular mechanism for GABA release that does not require VGAT also exists (i.e. via LRRC8 channels (Lutter et al., 2017)). Another important evidence for endogenous GABA tone is the *in vivo* studies that demonstrated that GABA reuptake inhibitor, NO711, produced analgesic effect when delivered to the DRG *in vivo* (Du et al., 2017; Obradovic et al., 2015). DRG-applied GABA_A_ antagonists, on the other hand, exacerbated peripherally-induced pain and even produced a pain-like behavior (Du et al., 2017). Hence, there must be ambient and efficacious levels of endogenous GABA in the DRG influencing action potential throughput.

A major causative factor of neuropathic pain is a loss of dorsal horn GABAergic inhibitory system (Moore et al., 2002). GABAergic progenitor cells implanted into adult spinal cord survive and integrate into the spinal inhibitory circuit (Braz et al., 2017; Etlin et al., 2016; Llewellyn-Smith et al., 2018), reducing neuropathic pain severity (Braz et al., 2012; Braz et al., 2015). We found the MGE cells transplanted into the DRG of adult mice also survive there and can release GABA (Fig. 3). Optogenetic stimulation of MGE-transplanted DRG significantly alleviated hypersensitivity to noxious stimuli induced by chronic inflammation (Fig. 3). Importantly, even without optogenetic stimulation, MGE cell transplantation accelerated the recovery from both inflammatory and neuropathic types of such hypersensitivity (Suppl. Fig. 8). This, together with observed higher efficiency of GABAergic filtering at the C, as compared to A fibers might have a therapeutic significance as it suggests that GABAergic system in DRG could be targeted for pain relief without significantly compromising other haptic sensations. In the spinal cord MGE cells maturate into interneurons and integrate into the existing spinal inhibitory system (Braz et al., 2012; Etlin et al., 2016). For the case of the DRG transplant, the direct analogy is unlikely as there are no ‘classical’ interneurons in DRG. The most straightforward explanation for the anti-nociceptive effect of MGE cells in DRG is a ‘GABA pump’ mechanism, whereby MGE cells simply release GABA into the extracellular space thus increasing the GABA tone. Recent years saw increasing success in using stem cells as a chronic pain therapy (Hwang et al., 2016; Yu et al., 2015) and GABAergic progenitor cells can be generated from human stem cells (Liu et al., 2013) and integrated into the pain pathways (Manion et al., 2020). Thus, targeting GABAergic system with the DRG-directed stem cell therapy or GABA-mimetics tailored to reduce their CNS permeability could open up avenues for analgesic strategies with reduced CNS side effects.

### Study limitations

It must be noted, that whilst our approach to matching spikes between SN and DR recording sites is largely accurate in simulations, it does have some real-world limitations that are important to consider. These include the firing rate of the bulk nerve recording and the conduction velocity of the slowest conducting fiber. The accuracy decreases slightly when the firing rate increases, and also with very slow conduction velocities. Some variability may arise from different electrode characteristics between the recording sites and signal to noise ratios that affect spike extraction in the first instance, in addition to variability in the unsupervised method of spike sorting. However, with these considerations in mind and ensuring a short distance between recording sites, it is perfectly feasible to temporally correlate spikes from peripheral and central nerves across dorsal root ganglia t-junctions.

In this study we provide for the first time quantitative measurements of stem axon lengths for both C- and A-fiber neurons, along with measurements of somatic diameter. But our modeling approach is limited by several critical membrane parameters. Mostly notably, all the membrane mechanisms (i.e. voltage-gated conductances) available in the literature are based on somatic measurements. The properties of axonal ion channels may be quite different in terms of their density, voltage-dependence, and kinetics. As such, what we present here are minimal models with uniform distributions of several key conductances along the neuronal membrane (voltage- and ligand-gated channels), as well as a uniform concentration of ions and extracellular GABA throughout the tissue. It is highly likely that there is much more variability within and between cells in vivo. Future simulations, incorporating more experimentally-determined parameters and physiological measurements, will be necessary to improve and validate our computational approach.

## MATERIALS AND METHODS

All animal experiments performed in Hebei Medical University were in accordance with the Animal Care and Ethical Committee of Hebei Medical University (approval number: IACUC-Hebmu-2020007). All animal work carried out at the University of Leeds was approved by the University of Leeds Animal Welfare Ethical Review Body (AWERB) and performed under UK Home Office License P40AD29D7 and in accordance with the regulations of the UK Animals (Scientific Procedures) Act 1986.

### *In vivo* recording of peripheral nerve and dorsal root activity

All surgical procedures were performed under deep anesthesia with an i.p. injection of pentobarbital sodium (60 - 80 mg/kg) in accordance with the Animal Care and Ethical Committee of Hebei Medical University under the International Association for the Study of Pain guidelines. In one set of experiments (fig. S4) pentobarbital was replaced by isoflurane (4% for induction, 2% for maintenance). Laminectomy was performed to expose right L5 DRG of adult male rat (Sprague-Dawley, 180-200 g) or L4 DRG of adult male C57BL/6J mice. Dorsal root (DR), spinal nerve (SN) and DRG were exposed by removal of both spinous and transverse processes of the vertebra bone; the DR and SN were then suspended (without transection) on the hooked stainless-steel recording electrodes connected to BL-420F biological data acquisition and analysis system (Chengdu Techman Software Co., Ltd. China). The wound was filled with paraffin in order to stabilize preparation. The right hindpaw was used for the injection of capsaicin (10 µM; 50 µl for rat; 20 µl for mouse) or Bradykinin (100 µM, 50 µl for rat; 20 µl for mouse), or the stimulation with hot water (~60°C), ice, air puffs (using bulb syringe), von Frey filaments (4 g for rat; 0.4 g for mouse) or needle prick (glass electrode). GABA (200 µM; 3 µl for rat; 2 µl for mouse) or Bicuculline (200 µM; 3 µl for rat; 2 µl for mouse) was accurately delivered to the surface of exposed DRG by micropipettor.

For the single unit recording in rats, dorsal root and spinal nerve were exposed and covered with liquid paraffin. A single axon bundle was teased away from dorsal root by a fine tweezer and placed on nerve fiber electrode for electrophysiological recording. The stimulus current pulses (2 - 5 mA) were delivered to the spinal nerve at 10/50/100 Hz. Conduction velocity of fiber was determined by dividing conduction distance by response latency. GABA (200 µM, 3 µl), Tetrodotoxin (TTX, 1 µM, 3 µl) or vehicle (3 μl saline) were accurately delivered to DRG by micropipettor. During the operation and recording, the animal body temperature was maintained at 38 °C with the Animal Temperature Controller pad (RWD Life Science Co. Ltd., China).

### Ventral root transection and ChAT immunohistochemistry

Laminectomy was performed to expose right L5 DRG of adult male rat (Sprague-Dawley, 180-200 g). Ventral root (VR) was exposed by removal of both spinous and transverse processes of the vertebra and transected (5mm of the nerve removed). The muscle and skin were sutured, area of the wound was disinfected with iodophor and the animals were transferred to a recovery cage. Two weeks after ventral root transection, the rats were subjected to electrophysiological recordings or sacrificed for immunohistochemical detection of ChaT-positive motor fibers. In the latter case, the spinal nerve sections of L5 DRGs were removed and submerged in Tissue-Tek O.C.T. (Sakura, Alphen aan den Rijn, The Netherlands), frozen, and sectioned (10 μm) using a freezing microtome (CM1950, Leica Microsystems). Sections were placed on microscope slides, and washed thrice with 0.01 M PBS (Sigma-Aldrich), fixed in 4% PFA (Biosharp) for 1 hour, and blocked for 2 hours with blocking buffer (3% donkey serum in 0.1 M PBS; Sigma-Aldrich). Primary antibodies (ChAT, Abcam, 1:400) were diluted in 0.3% Triton X-100/PBS buffer before overnight incubation at 4°C. The following day, sections received a further 3 washes in PBS before incubation with secondary antibodies (Alexa Fluor 488, 1:500) for 2 hours at room temperature. Sections were washed with PBS 3 times and Sealed with cover glass. Staining was visualized using a confocal fluorescent microscope (TCS SP5 II, Leica).

### Spike sorting

Electrophysiological recordings of both the spinal nerve and dorsal root were imported into Python and high pass filtered at 60 Hz using a digital Butterworth filter from the SciPy module (Virtanen et al., 2020). Extracellular spike times and waveforms were extracted using an absolute median deviation of between 5 and 6 from the median of the signal. Extracted spike waveforms were sorted using the WaveClus program (Chaure et al., 2018) in Matlab to define individual neuronal units underlying the extracellular signal. Matching of spikes in the dorsal root to an origin spike in the spinal nerve was achieved by finding the minimum latency (within a tolerance window of the slowest theoretical fibre conduction velocity of 0.1 m/s) between spikes in the spinal nerve and dorsal root. The overall approach to spike sorting was as follows. (i) The SN spike waveforms were sorted to group SN spikes into ‘firing units/clusters’. (ii) For every spike in DR we tried to identify an origin spike in the SN, based on minimal latency with a cut-off of 150 ms (calculated from the slowest expected conduction velocity of C fibres of 0.1 m/s over the estimated distance between two recording electrodes of 1.5 cm). (iii) The spike identity, defined by the SN spike waveforms, was extended to the DR spikes after matching. (iv) For each firing unit/cluster, the propagation success for each firing unit was quantified based on how many SN spikes in that firing unit had a matched DR spike (Number of SN spikes with DR match/Total SN spikes) x 100.

To validate this approach we used a computer-generated, Poisson randomly generated spike train to simulate an SN recording (Suppl. Fig. 5). The DR spike train was produced by firstly randomly assigning ‘spikes’ in the simulated SN spike train a ‘fiber type’ from Aα, Aβ, Aδ and C. To introduce randomness and variability, each SN ‘spike’ was randomly assigned a conduction velocity from a normal distribution of the expected mean conduction velocity for the assigned fibre type. The randomly generated conduction velocity for each SN spike dictated the latency with which an associated spike should be placed in the simulated DR spike train as if it was recorded 1.5cm away from the simulated spinal nerve spike train (as per our *in vivo* recording conditions). To evaluate the performance of spike-matching during t-junction filtering, spikes in the DR were randomly deleted to simulate different degrees of filtering. Accuracy was defined as the percentage of correct matches from total ‘ground truth’ matches (Suppl. Fig. 5 B). We hypothesized that this latency-based spike-matching method would decrease in accuracy as the firing rate increased as this would increase the likelihood of spikes, travelling at different velocities, overlapping in the DR spike train. In a similar vein, we also hypothesized that the accuracy would be determined by the slowest conducting fibre as this again would increase the likelihood of faster spikes overlapping with the slower spikes. Both predications were indeed confirmed. We found that the accuracy of our method was reduced to 80% if a very slow conducting mean C fibre velocity was used to randomly shift dorsal root spike latencies and also at higher spinal nerve firing rates (Suppl. Fig. 5B). A false-positive in this case was more likely to be a faster spike overlapping with a slower spike i.e. incorrectly finding a match for a slow spinal nerve spike. We simulated spinal nerve firing rates up to 100Hz but none of the in vivo spike sorted recordings approached this firing rate giving us confidence that little overlap was occurring and that our method was accurate (>80%) at our observed firing rates. We also independently adjusted the mean conduction velocities of other fibre types and found no effect on accuracy. To further increase randomness in our assessment of this method, each simulation experiment (where a fibre type mean conduction velocity was adjusted, at varying spinal nerve firing rates and degrees of random deletion) was repeated five times with new randomly generated Poisson spike trains, with new randomly assigned conduction velocities. Source data and code for spike sorting analysis is available at GitHub (https://github.com/pnm4sfix/SpikePropagation).

### MGE cells transplantation

The female VGAT-ChR2-EYFP mice (Jackson Laboratory) with embryos between E12.5 and E13.5 were anesthetized with an intraperitoneal injection of pentobarbital sodium (60-80 mg/kg). All the embryos were removed via abdominal incision and placed into a 10 cm Petri dish containing ice-cold HBSS. The mice were humanely sacrificed. The gestational sac of each embryo was removed using forceps under the stereo microscope (SZX7, Olympus). The embryonic brain was extracted and cut along the sagittal plane to separate two hemispheres. Medial ganglionic eminence (MGE) on each hemisphere was then removed with a scalpel. MGE tissue was put in a 1.5 ml collection tube containing 500 μl DMEM/10% FBS (Sigma) and triturated into a single cell suspension as described(Vogt et al., 2015). The final cell density in the suspension of embryonic stem cells was measured with hemocytometer. Adult (5-6 weeks) male C57BL/6J mice were anesthetized with an intraperitoneal injection of pentobarbital sodium (60 - 80 mg/kg). L4 DRG was exposed by removal of both spinous and transverse processes of the vertebra bone. The microinjector (Hamilton Co.) loaded with a suspension of MGE cells (3 µl; ~1×10^7^/ml) was inserted into the ganglion to a depth of 200 μm from the exposed surface. The cell suspension was injected slowly, and the needle was removed 3 minutes after the injection. The muscles overlying the spinal cord were loosely sutured together, and the wound was closed. Animals developing signs of distress were humanely sacrificed. In order to verify that the DRGs were transplanted with MGE cells successfully, the L4 DRGs were excised, submerged in Tissue-Tek O.C.T. (Sakura, Alphen aan den Rijn, The Netherlands), frozen, and sectioned (10 µm) using a freezing microtome (CM1950, Leica Microsystems). Slices were then analyzed for the EYFP fluorescence using confocal microscopy (TCS SP5 II, Leica Microsystems).

### Chronic pain models

Chronic constriction injury (CCI) was performed as described previously (Du et al., 2017). Briefly, rats were anesthetized with an i.p. injection of sodium pentobarbital (60-80 mg/kg). The right hind leg was shaved and cleaned using 70% ethanol. The sciatic nerve was exposed by blunt dissection at the mid-thigh level, proximal to the sciatic trifurcation. Four nonabsorbable sterile surgical sutures (4-0 chromic gut; packaged in isopropyl alcohol and soaked in saline for 10 min prior to application) were loosely tied around the sciatic nerve with an approximately 1.0-to-1.5-mm interval between the knots. The skin was sutured, and the animal was transferred to a recovery cage. To induce chronic inflammatory pain, CFA (20 μl) was injected into the plantar surface of the right hind paw of the mice.

### Behavioral tests

Mechanical withdrawal threshold was measured by a set of von Frey filaments (Stoelting Co, Chicago, IL, USA) with a calibrated range of bending force (0.16, 0.4, 0.6, 1, 1.4, 2g). Each mouse was placed into a plastic cage with a wire mesh bottom. A single filament was applied perpendicularly to the plantar surface of hind paw for five times with an interval of 5s. Positive response was defined as at least three clear withdrawal responses out of five applications. Filaments were applied in an up-and-down order according to a negative or positive response to determine the hind paw withdrawal threshold. Thermal withdrawal latency was tested by a radiant heat lamp source (PL-200, Taimeng Co, Chengdu, China). The intensity of the radiant heat source was maintained at 10±0.1%. Mice were placed individually into Plexiglas cubicles placed on a transparent glass surface. The light beam from radiant heat lamp, located below the glass, was directed at the plantar surface of hindpaw. The time was recorded from the onset of radiant heat stimulation to withdrawal of the hindpaw. Three trials with an interval of 5 minutes were made for each mouse, and scores from three trials were averaged.

### *In vivo* optogenetic stimulation

Adult male C57BL/6J mice were L4 DRG transplanted with MGE cells from VGAT-ChR2-EYFP mice 3-4 weeks before experiments. Recordings of DR and SN activity was performed as described above in combination with laser stimulation (473 nm, 3 mW, 30 Hz for 10 seconds with 20-second interval) of DRG using an MLL-FN-473-50 unit (Changchun New Industries Optoelectronics Technology Co., Ltd.) controlled by a pulsing set (S48 Stimulator, Grass Technologies, An Astro-Med, Inc. Product Group). In the behavioral tests on freely moving animals, a stainless steel cannula guide (RWD Life Science Co. Ltd., China; diameter 0.64 mm) was implanted inot the L4 DRG, the cannula was firmly fixed in place with dental cement, and the optical fiber (RWD Life Science Co. Ltd., China; diameter 0.2 mm, length 1 m) was inserted through the guide; a more detailed description is provided in (Du et al., 2017).

### Patch clamp recording from DRG neurons

DRG dissection and recording were performed as described previously (Liu et al., 2010). Briefly, L4 DRG transplanted with MGE cells 4 weeks in advance was carefully removed and digested with a mixture of 0.4 mg/ml trypsin (Sigma) and 1.0 mg/ml type-A collagenase (Sigma) for 45 min at 37 °C. The intact ganglia were then incubated in ACSF (artificial cerebrospinal fluid) oxygenated with 95% O2 and 5% CO2 at 28 °C for at least 1 h before transferring them to the recording chamber. DRG were visualized with a 40X water-immersion objective using a microscope (BX51WI; Olympus, Tokyo, Japan) equipped with infrared differential interference contrast optics. Whole-cell current recording was acquired with an Axon700B amplifier (Molecular Devices Corporation, Sunnyvale, CA, USA) and pClamp 10.0 software (Axon Instruments); recordings were sampled at 5 kHz. Patch pipettes (4-7 MΩ) were pulled from borosilicate glass capillaries on P-97 puller (Sutter Instruments, USA). The series resistance was 10-20 MΩ. Continuous voltage-clamp recording was performed via holding potential of −60mV. The ACSF contained (in mM): 124 NaCl, 2.5 KCl, 1.2

NaH2PO4, 1.0 MgCl2, 2.0 CaCl2, 25 NaHCO3, and 10 Glucose. The pipette solution contained (in mM): 140 KCl, 2 MgCl2, 10 Hepes, 2 Mg-ATP, pH 7.4. Osmolarity was adjusted to 290-300 mOsm. For optical stimulation of MGE cells derived from VGAT-ChR2-EYFP mice, a 473-nm blue light (3 mW) was elicited using the same device as for in vivo stimulation.

### VGAT antibody uptake

DRG neurons were dissociated and cultured as described previously(Liu et al., 2010). DRG neurons were incubated for 15 min in either normal or ‘high K^+^’ extracellular (EC) solution supplemented with either C-terminal (luminal) N-terminal (cytosolic) VGAT antibodies. Normal EC solution contains (in mM): 144 NaCl, 5.8 KCl, 1.3 CaCl_2_, 5.6 D-glucose, 0.7 NaH_2_PO4, 0.9 MgCl_2_ and 10 HEPES (all from Sigma). In high K^+^ EC solution NaCl concentration was lowered to 49.8 mM and KCl concentration was raised to 100 mM. Ca^2+^-free EC solution was also used; in this solution CaCl_2_ was omitted. After incubation cell cultures were washed 3 times with PBS and fixed using 4% paraformaldehyde, followed by permeabilization with 0.05% tween 20 and 0.25% triton-X 100 (with donkey serum) for 1 hr. Cells were then labelled with secondary antibody, washed three times with PBS, mounted on coverslips and imaged using Zeiss LSM880 confocal microscope. The following antibodies were used: VGAT C-terminal antibody (rabbit polyclonal #AB-N44, Advance Targeting System; 1:200); VGAT N-terminal antibody (rabbit polyclonal 131002, Invitrogen, Eugene, Oregon, USA; 1:1000); secondary antibody: alexafluor donkey anti-rabbit 488 (Invitrogen, Eugene, Oregon, USA; 1:1000). Cells were deemed positively stained if the mean fluorescence intensity of the somatic area was at least two times of the background fluorescence intensity.

### Iodide imaging

HEK293 cells were co-transfected with cDNA encoding human α1, β2 and γ2 subunits of GABA_A_ receptors (gift of David Weiss, Department of Physiology, University of Texas Health Science Center, San Antonio, Texas, USA) together with the halide-sensitive EYFP mutant (H148Q/I152L; EYFP-QL) using FuGENE® HD transfection reagent. Transfected cells were co-cultured for 24 hrs with DRG neurons isolated as described above (see also (Du et al., 2017)). Extracellular solution consisted of (mM): NaCl (160); KCl (2.5); MgCl_2_ (1); CaCl_2_ (2); HEPES (10) and Glucose (10); pH adjusted to 7.4 with NaOH (All from Sigma). I^-^ containing solution was produced by equimolar substitution of 5 mM NaCl with NaI. I^-^ imaging was performed using Nikon TE-2000 E Swept Field Confocal microscope using 488nm argon laser as excitation light source. Images were recorded and analyzed using Nikon Elements software.

### Dorsal root ganglia clearing, staining and morphometry

DRGs were extracted from the lumbar spinal column of euthanized adult Wistar rats (150-250g) and immersion fixed in 4% paraformaldehyde at 4 °C overnight. Tissue clearing of the ganglia was performed using the iDISCO+ protocol (Renier et al., 2016). Briefly, DRG samples were pre-treated and permeabilized before blocking with 3% donkey serum. Primary antibodies raised against Neurofilament-200 (NF-200, 1:500, Sigma-Aldrich, N5389) and peripherin (Abcam, 1:250, ab39374) were used as markers for myelinated neurons and nociceptors, respectively. DRGs were incubated in primary antibodies for five days at 37 °C, washed for 24 hours and incubated in secondary antibodies (donkey anti-mouse 555, 1:1000, donkey anti-chicken 488, 1:1000, Invitrogen, A31570, A21202) for four days at 37 °C. Following final washes DRGs were embedded in 1% agarose cubes, sequentially dehydrated in a methanol/H_2_0 series (20%, 40%, 60%, 80%, 100%, 100%) and stored overnight in 100% methanol at room temperature. Following methanol dehydration, samples were incubated for three hours in 33% methanol/66% dichloromethane (DCM, Sigma-Aldrich) solution. To complete the tissue clearing, this was followed by two 15-min incubations in 100% DCM and final storage in DiBenzyl Ether (DBE, Sigma-Aldrich).

Cleared DRG samples were imaged using the 20X objective of an Ultramicroscope II (LaVision BioTech) resulting in an axial resolution of 0.35 × 0.35 μm. The numerical aperture of the light sheet was 0.15 and resulted in a light sheet focal thickness of 4 μm. Dynamic horizontal focusing was utilized to ensure a homogenous point spread function across the field of view. Images were analyzed using the simple neurite tracer in FIJI (Longair et al., 2011; Schindelin et al., 2012). Using this semi-automatic method of tracing, stem axons were measured from origin at peripherin and NF-200-positive somata through the image stack until clear bifurcation or in the case of NF-200-positive cells, loss of a clear signal.

### Computer modeling

All simulations were performed using NEURON (Hines and Carnevale, 2001) on an Intel-based Macintosh computer (http://neuron.yale.edu) and analyzed using Python scripts. Representative code is available at GitHub (https://github.com/dbjaffe67/DRGsims). A simplified model of a portion of an Aδ neuron within the DRG was constructed of a spinal nerve (SN) and dorsal root (DR) axon of 21 mm length of 4 and 3 μm diameter, respectively, of alternating internodal (150 μm length) and nodal (1.5 μm) segments (3 μm diameter) (Hoheisel and Mense, 1987). A stem axon was connected to the t-junction node midway along the SN/DR axis. Stem axon internode distance was 100 μm with nodes of 1.5 μm, both with diameters ranging from 1-3 μm. An unmyelinated axon initial segment (AIS) (Nascimento et al., 2018) of 50 μm and 2 μm diameter was attached to the last stem node and the soma (40 μm diameter (Villiere and McLachlan, 1996)). All internodal/myelin wrapped segments had a specific membrane conductance (G_m_) of 10 μS/cm^2^ and a specific membrane capacitance (C_m_) of 0.01 μF/cm^2^, while for the nodes, AIS, and soma the values for G_m_ and C_m_ were 200 μS/cm^2^ and 1 μF/cm^2^, respectively. Soma and nodes contained a TTX-sensitive voltage-gated Na^+^ conductance (G_Na_) reflecting a mix of Na_V_1.1, 1.6, and 1.7 channels (Herzog et al., 2001). Spike repolarization in nodes was achieved solely by passive leak conductance (Chiu et al., 1979), while both the AIS and soma also contained a delayed rectifier conductance (5 mS/cm^2^). A chloride conductance (G_Cl_) with E_Cl_ = −40 mV (Liu et al., 2010) was added to model tonic GABA_A_ receptor activation in various compartments (i.e. soma and/or axons). Resting potential was set in all compartments to −60 mV (Du et al., 2014; Villiere and McLachlan, 1996) and used to calculate the resting leak equilibrium potential from the sum of steady-state resting currents. Action potentials were initiated by injecting suprathreshold current (1 ms duration) into the most distal portion of the spinal nerve axon. Morphology for an unmyelinated C fiber neuron, used for comparison with the A-neuron, was based on previously published models (Du et al., 2017) constructed with SN/DR axons of 5.5 mm each with diameters of 0.8 and 0.4 μm, respectively. The stem axon was connected at the midpoint with a length ranging from 25-150 μm and diameters 1-2 μm. The same TTX-sensitive voltage-gated Na^+^ conductance was added to all compartments and spike repolarization was achieved by a delayed rectifier K^+^ conductance (Sheets et al., 2007). Specific membrane resistivity for all compartments was 10,000 Ωcm^2^, intracellular resistance 100 Ωcm, and capacitance of 1 μF/cm^2^

### Statistics

Sample size estimations were made using NC3Rs Experimental Design Assistant (https://eda.nc3rs.org.uk/). Animals were allocated to the groups randomly. In the instances where blinding was appropriate (e.g. in vivo optogenetic stimulation of implanted MGE cells) the operator was unaware of specific surgery allocations. All data are given as mean ± SEM. Paired or unpaired t-test was used to compare the mean of two groups when the data were normally distributed. Multiple groups were compared using ANOVA (one-, two- or three-factor) or repeated-measures ANOVA, depending on experimental setup; Bonferroni post-hoc test was used for comparison between groups; for data failing normality test Kruskal-Wallis ANOVA with Mann-Whitney post-hoc test was used. Statistical analyses were performed using IBM SPSS Statistics 21, GraphPad Prism or Origin. Statistical parameters are given in the figure legends.

## Supplemental material

Current version of the manuscript includes 13 Supplemental Figures (Suppl. Fig. 1-13) and 2 Supplemental Movies (Movie S1, S2).

## Acknowledgments

We thank Prof Jim Deuchars and Dr Ronaldo Ichiyama (University of Leeds) for advice on experiments and helpful suggestions. We also thank Dr Junling Xing (Fourth Military Medical University) for help with setting up single unit recordings.

## Author Contributions

N.G. and X.D. designed and directed the study; H.H., R.R., C.W., C.L., S.S., P.M., V.L., F.J., J.S. and N.X. conducted the experiments and analyzed data; D.B.J performed computational modeling; N.G., X.D. and H.H. wrote the paper, assisted by all the coauthors.

