## Supplemental figures and legends for "Dorsal root ganglia control nociceptive input to the central nervous system"

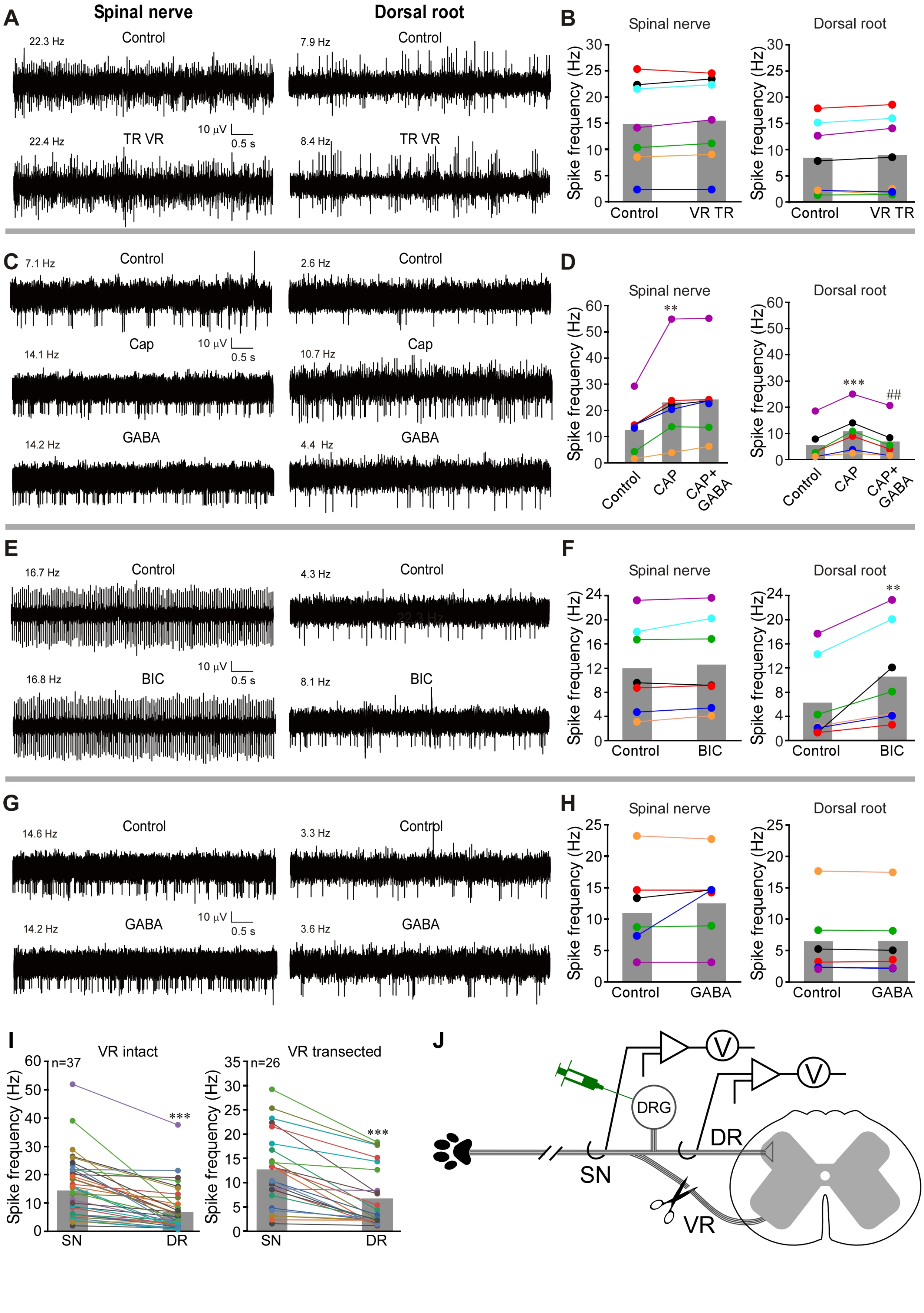


Supplemental Figure 1

**Supplemental Figure 1. Ventral root transection does not affect filtering at the DRG.** (**A**) VR transection (schematized in the panel J) does not affect spontaneous activity in the SN or DR. (**B**) Summary of experiments exemplified in A. Two-factor (nerve site, VR transection) repeated measures ANOVA: main effect associated with nerve site [$F(1,12)=12.7$; $p<0.05$). (**C**) After the VR transection a baseline was recorded (control) and Capsaicin (CAP, 10 µM, 50 µl) was injected into the hindpaw. CAP increased firing frequency in both SN and DR branches of the nerve (middle traces, as compared to basal activity shown in the upper traces). Application of GABA (200 µM, 3 µl) to DRG reduced CAP-induced firing frequency in DR but not SN (bottom traces). (**D**) Summary of the panel C. Two-factor (nerve site, drug application) repeated measures ANOVA: main effects associated with nerve site [$F(1,10)=12.2$; $p<0.05$] and drug application [$F(2,9)=6.5$; $p<0.05$]; significant interaction between nerve site and drug application [$F(2,9)=41.3$; $p<0.01$]. Bonferroni post-hoc test: **,***significant difference from control ($p<0.01$, $p<0.001$); $^{\#\#}$significant difference from CAP ($p<0.01$). (**E**) After the VR transection a baseline was recorded (control) and GABAA antagonist bicuculline (BIC, 200 µM, 3 µl) was applied to DRG; hindpaw was not stimulated. (**F**) Summary for panel E. Two-factor repeated measures ANOVA: main effects associated with nerve site [$F(1,12)=8.2$; $p<0.05$), drug application [$F(1,12)=18.2$; $p<0.01$), significant interaction between nerve site and drug application [$F(1,12)= 8.1$; $p<0.05$]. Bonferroni post-hoc test: **significant difference from control ($p<0.01$). (**G**) Experiments similar to these shown in (E, F) but GABA was applied instead of BIC. (**H**) Summary for panel G. Two-factor repeated measures ANOVA: main effect associated with nerve site [$F(1,10)=11.9$; $p<0.05$). (**J**) Scatter plot comparison of the firing basal (tonic) firing rates in paired SN-DR recordings with VR intact (left, data from the Fig. 1; Paired t-test: $t(25)=7.2$, $p<0.001$) and VR transected (right; Paired t-test: $t(36)=6.8$, $p<0.001$). (**J**) Schematic of the preparation. The overall procedure is similar to that in Fig.1 but ventral root (VR) is transected before (or during) the recording.

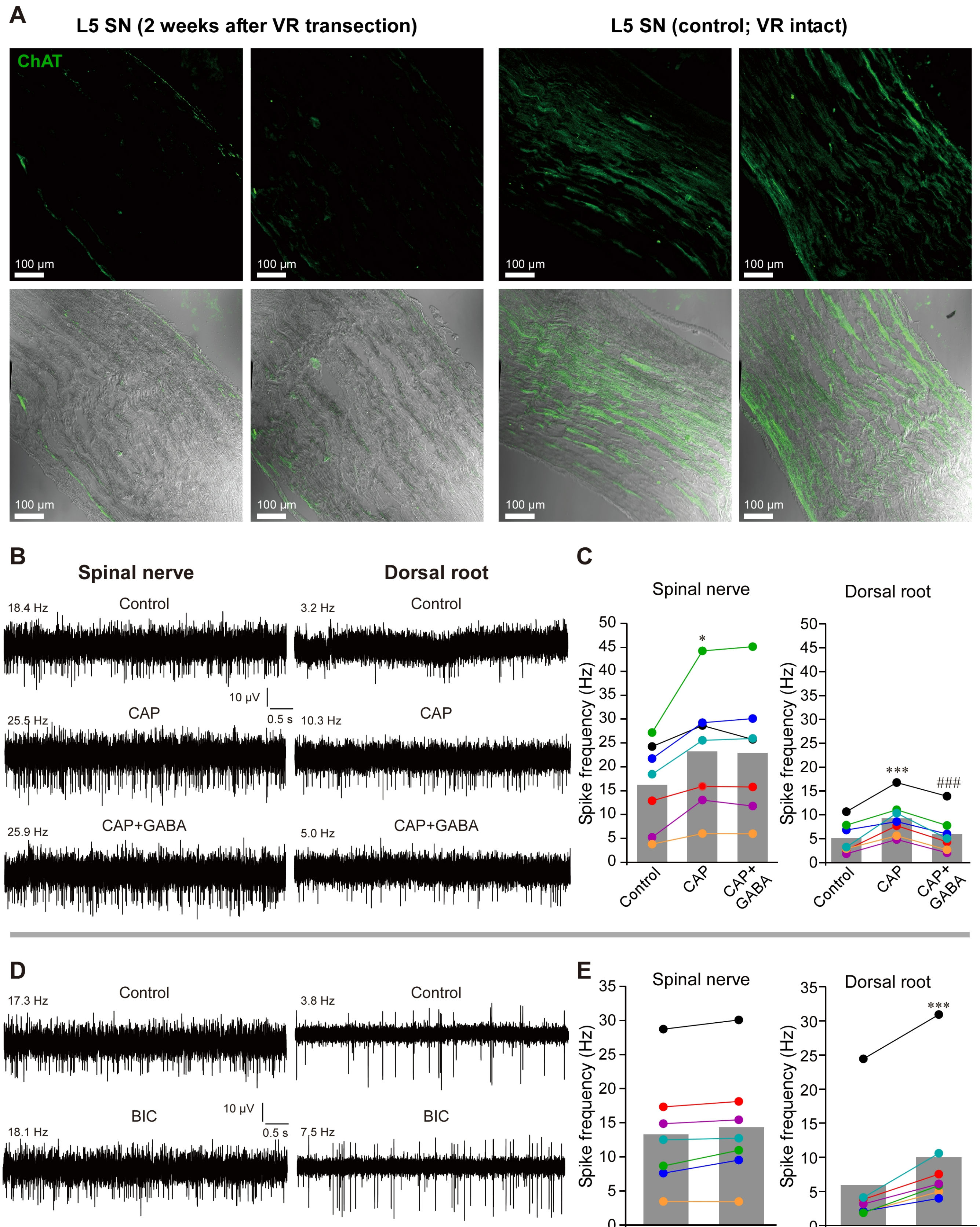


**Supplemental Figure 2**

**Supplemental Figure 2. Efferent fiber elimination does not affect filtering at the DRG.** The ventral root was surgically transected as in Suppl. Fig. 1. The animals were left to recover (see Suppl. Methods) and further experiments were done 2 weeks after the transection. (**A**) Immunostaining of the L5 spinal nerve of control (right) and VR-transected animals for Choline Acetyltransferase (ChAT). (**B**) 2 weeks after the VR transection a baseline was recorded (control) and Capsaicin (CAP, 10 µM, 50 µl) was injected into the hindpaw. CAP increased firing frequency in both SN and DR branches of the nerve (middle traces, as compared to basal activity shown in the upper traces). Application of GABA (200 µM, 3 µl) to DRG reduced CAP-induced firing frequency in DR but not SN (bottom traces). (**C**) Summary for panel B. Two-factor (nerve site, drug application) repeated measures ANOVA: main effects associated with nerve site [$F(1,12)=15.8$; $p<0.01$] and drug application [$F(2,11)=34.8$; $p<0.01$]; significant interaction between nerve site and drug application [$F(2,11)=7.7$; $p<0.05$]. Bonferroni post-hoc test: *,***significant difference from control ($p<0.05$, $p<0.001$); ##significant difference from CAP ($p<0.01$). (**D**) $GABA_A$ antagonist bicuculline (BIC, 200 µM, 3 µl) was applied to DRG; hindpaw was not stimulated. (**E**) Summary for panel D. Two-factor repeated measures ANOVA: main effects associated with nerve site [$F(1,12)=12.9$; $p<0.05$), drug application [$F(1,12)=52.9$; $p<0.001$), significant interaction between nerve site and drug application [$F(1,12)=14.8$; $p<0.01$]. Bonferroni post-hoc test: ***significant difference from control ($p<0.001$).

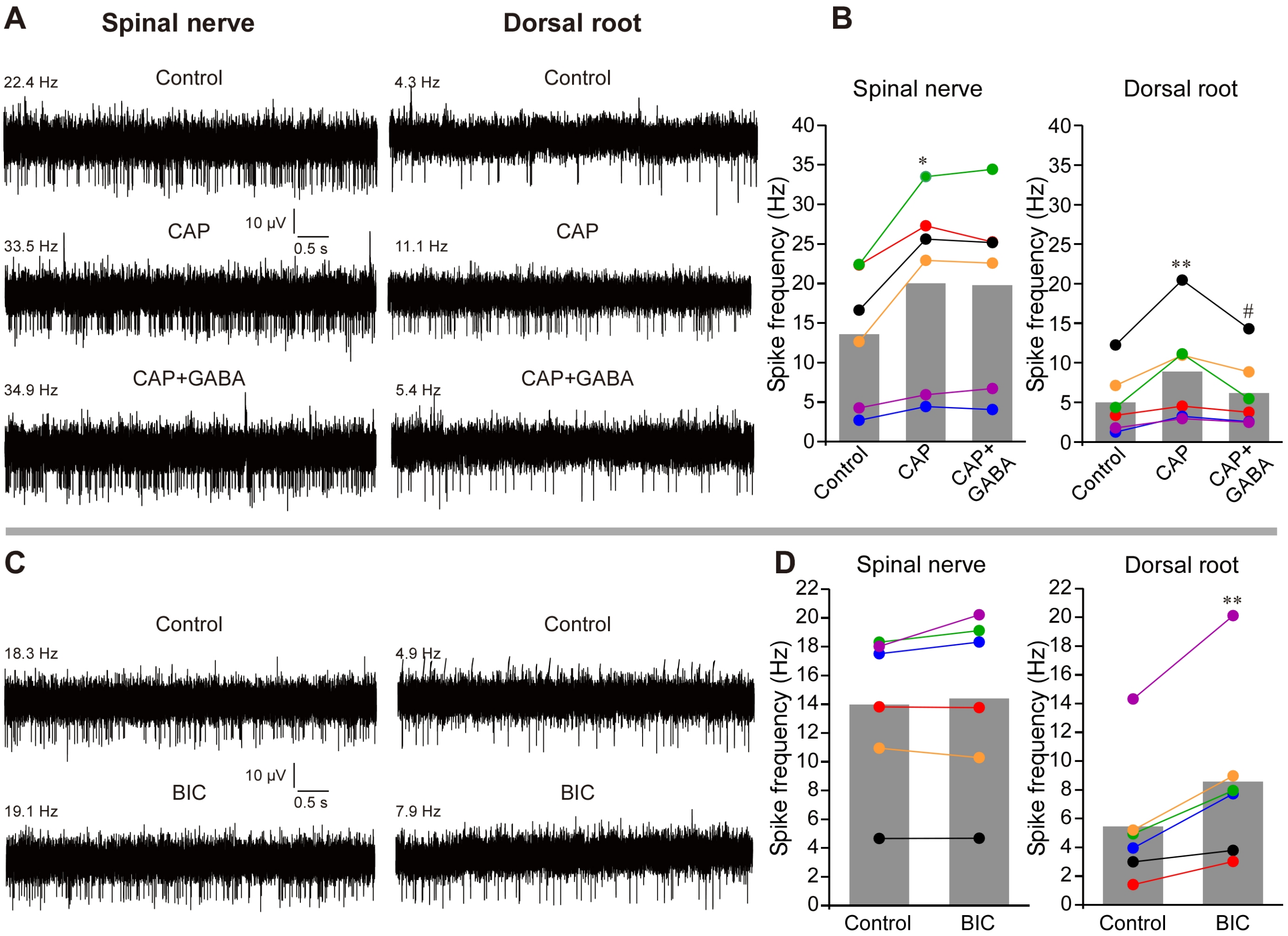


**Supplemental Figure 3**

**Supplemental Figure 3. DRG filtering in the female rats.** The recordings similar to that shown in Fig.1 but performed in female rats. (**A**) After baseline was recorded (control), Capsaicin (CAP, 10 μM, 50 μl) was injected into the hindpaw. Application of GABA (200 μM, 3 μl) to DRG reduced CAP-induced firing frequency in DR but not SN (bottom traces). (**B**) Summary for panel A. Two-factor (nerve site, drug application) repeated measures ANOVA: main effect associated with nerve site [F(1,10)=8.7; $p<0.05$]. Bonferroni post-hoc test: *,**significant difference from control ($p<0.05$, $p<0.01$); #significant difference from CAP ($p<0.05$). (**C**) $GABA_A$ antagonist bicuculline (BIC, 200 μM, 3 μl) was applied to DRG; hindpaw was not stimulated. (**D**) Summary for panel C. Two-factor repeated measures ANOVA: main effects associated with nerve site [F(1,10)=10.4; $p<0.05$), drug application [F(1,10)=12.2; $p<0.05$), significant interaction between nerve site and drug application [F(1,10)=23.1; $p<0.01$]. Bonferroni post-hoc test: **significant difference from control ($p<0.01$).

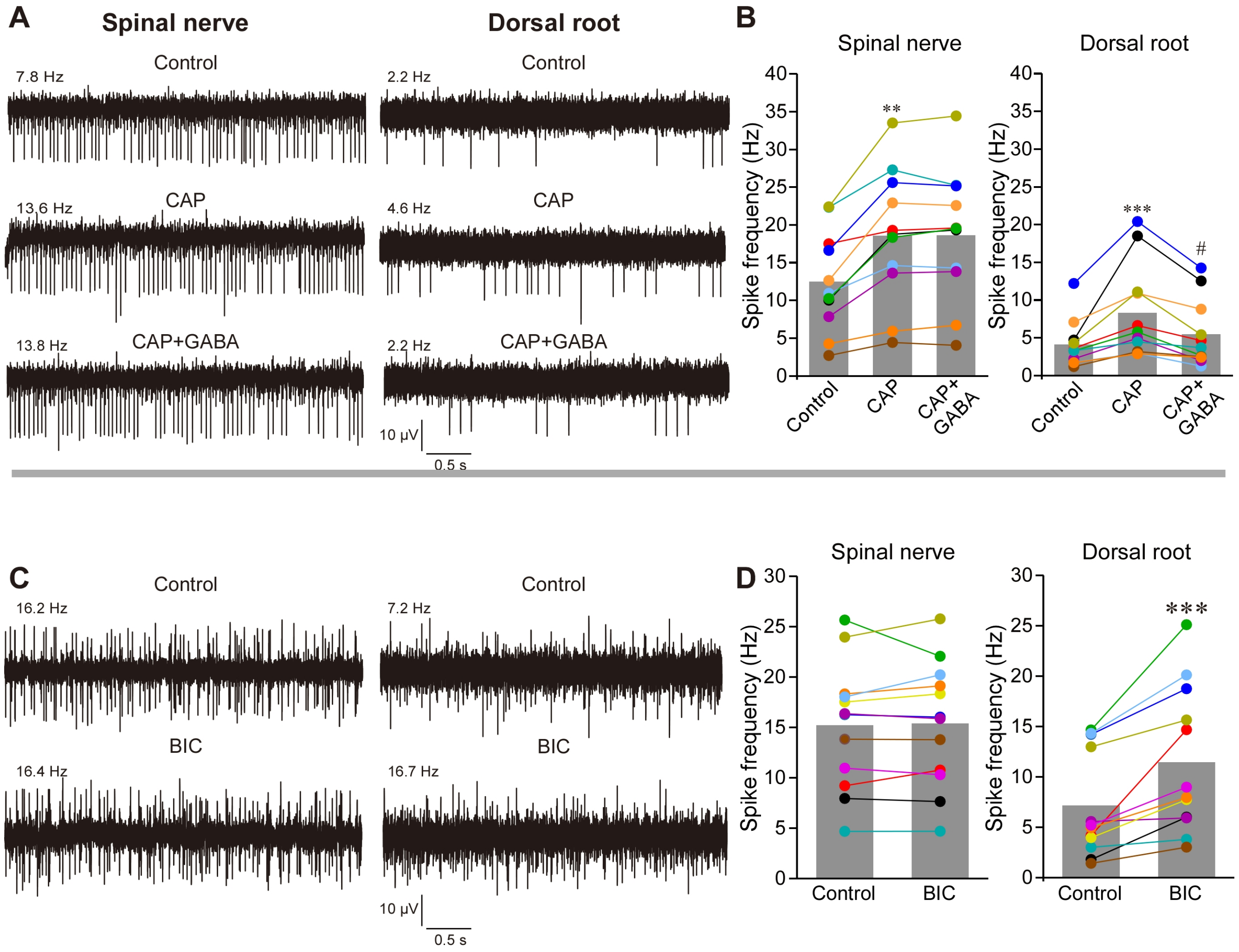


**Supplemental Figure 4**

**Supplemental Figure 4. DRG filtering as recorded in isoflurane anesthetized rats.** The recordings similar to that shown in Fig.1 but performed in male rats anesthetized with isoflurane. (**A**) After baseline was recorded (control), Capsaicin (CAP, 10 µM, 50 µl) was injected into the hindpaw. Application of GABA (200 µM, 3 µl) to DRG reduced CAP-induced firing frequency in DR but not SN (bottom traces). (**B**) Summary for panlel A. Two-factor (nerve site, drug application) repeated measures ANOVA: main effects associated with nerve site [$F(1,20)=25.5$; $p<0.001$] and drug application [$F(2,19)=11.8$; $p<0.01$]; significant interaction between nerve site and drug application [$F(2,19)=9.4$; $p<0.01$]. Bonferroni post-hoc test: **,***significant difference from control ($p<0.01$, $p<0.001$); #significant difference from CAP ($p<0.05$). (**C**) $GABA_A$ antagonist bicuculline (BIC, 200 µM, 3 µl) was applied to DRG; hindpaw was not stimulated. (**D**) Summary for panel C. Two-factor repeated measures ANOVA: main effects associated with nerve site [$F(1,22)=17.4$; $p<0.01$), drug application [$F(1,22)=21.4$; $p<0.001$), significant interaction between nerve site and drug application [$F(1,22)=13.6$; $p<0.01$]. Bonferroni post-hoc test: ***significant difference from control ($p<0.001$).

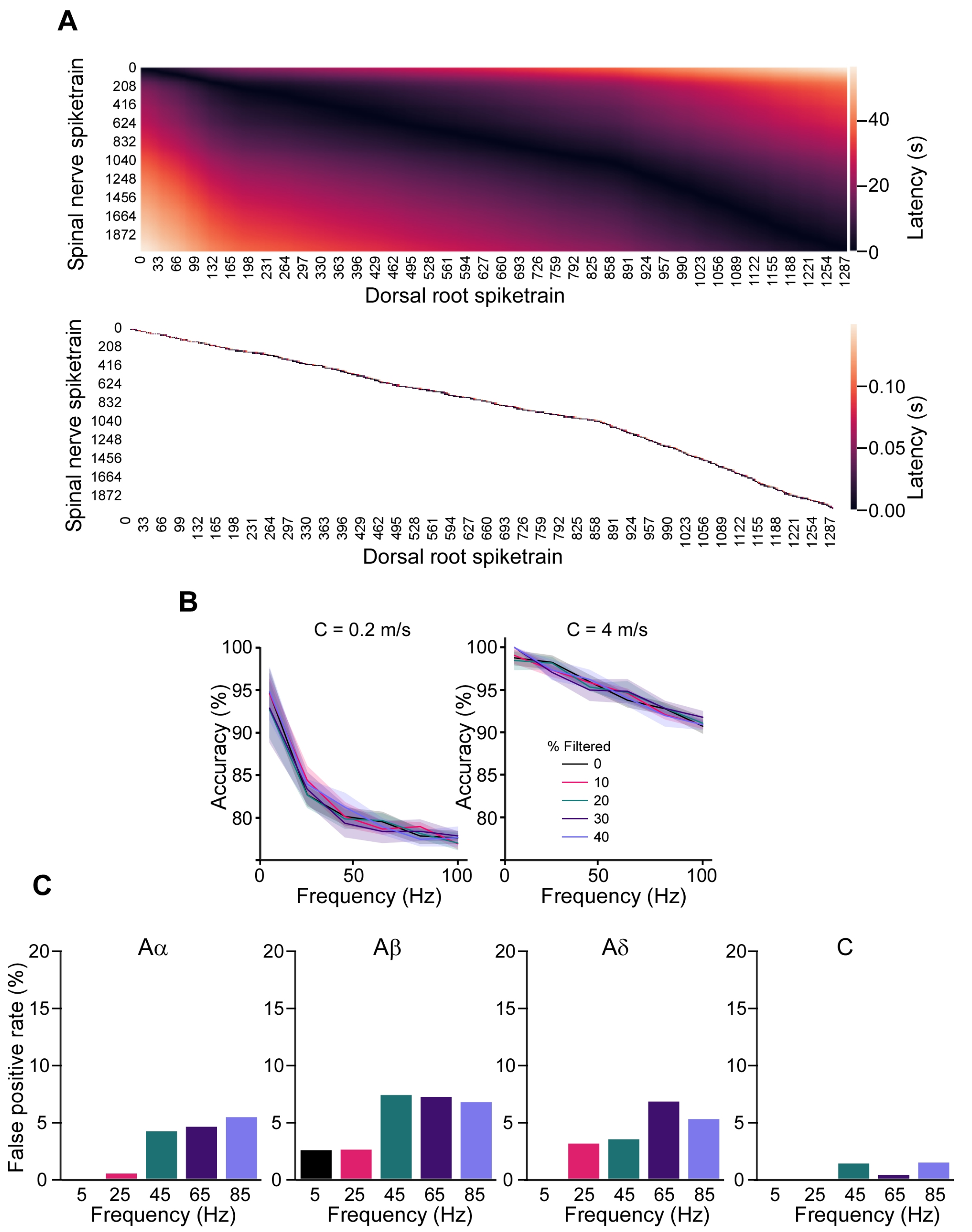


**Supplemental Figure 5**

**Supplemental Figure 5. Additional spike analyses.** (**A**) Top: the latency between SN and DR spikes was calculated; darker colours represent shorter latencies and hotter colours represent longer latencies. Bottom: each spike in the dorsal root was paired with a spike in the spinal nerve based on the minimum latency within a short time window. The end of the time window was defined by an estimation of the slowest conduction velocity of C-fibres; matching temporally uncorrelated spikes during gaps in spike activity was avoided. Using this method, the minimum latency defined the spinal nerve origin of a dorsal root spike. (**B**) The accuracy of latency-based spike matching at different firing frequencies in simulated poisson-generated spike trains. Filtering was modelled by random deletion of a percentage of DR spikes. Spikes were randomly assigned velocities which were varied around mean A and C fibre velocities. Spike matching accuracy was inversely proportional to firing frequency and was highly dependent on the velocity of the slowest fibre type, C fibre. (**C**) False positive rates (FPR) for fibre types at different simulated spike frequencies. The false positive rate was relatively low across fibre types but faster conducting fibre types were more likely to have a higher FPR at higher spike train frequencies.

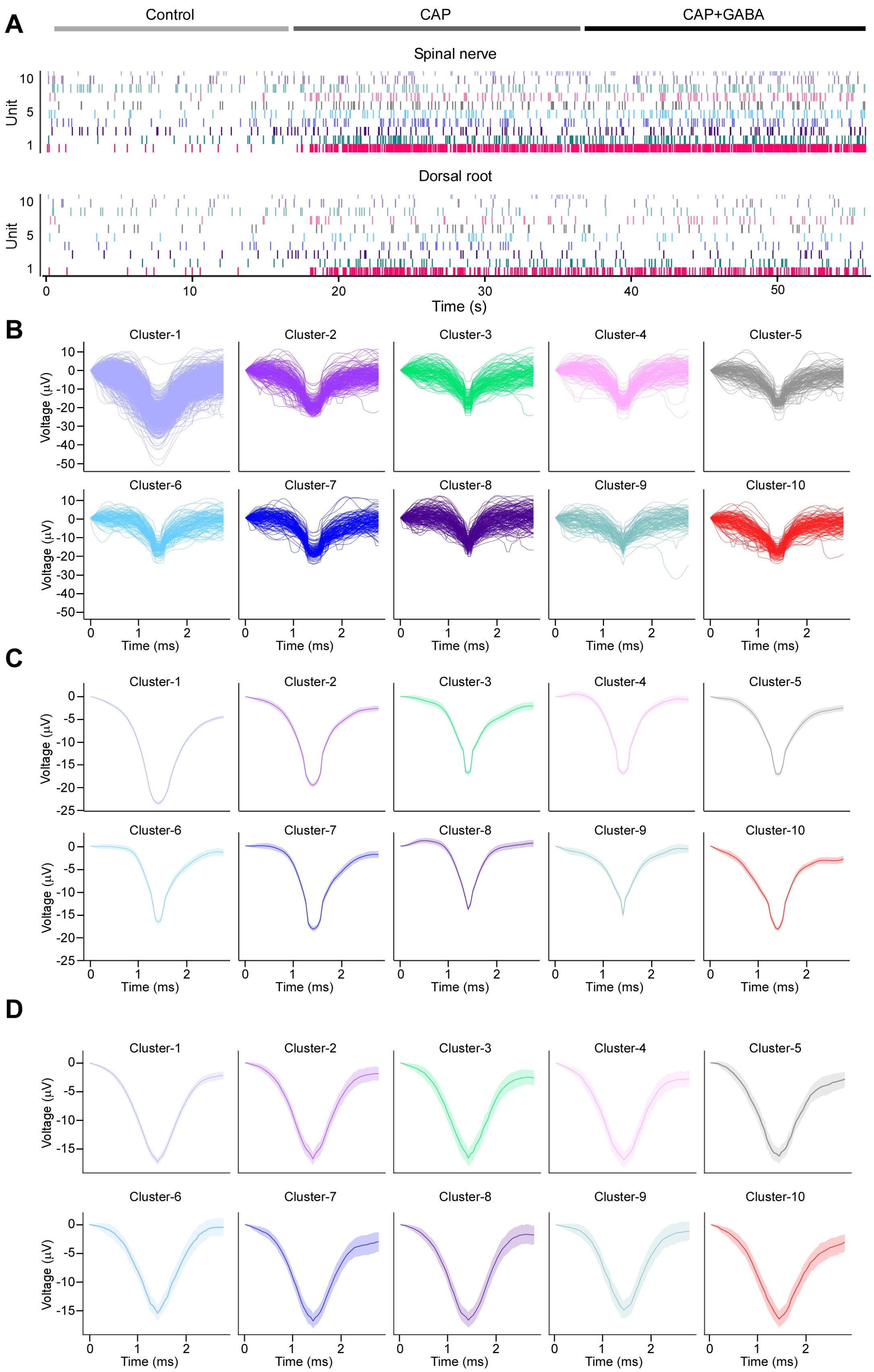


**Supplemental Figure 6**

**Supplemental Figure 6. Additional spike analyses.** (**A**) Raster plot for each clustered waveform (denoted as Unit) under control, CAP and CAP+GABA conditions after matching the DR spikes with these in the SN. (**B**) Individual waveforms of spike sorted units (cluster) in the SN. (**C**) Average waveforms of spike sorted units (clusters) in the SN. (**D**) Average waveforms of matched units in the DR.

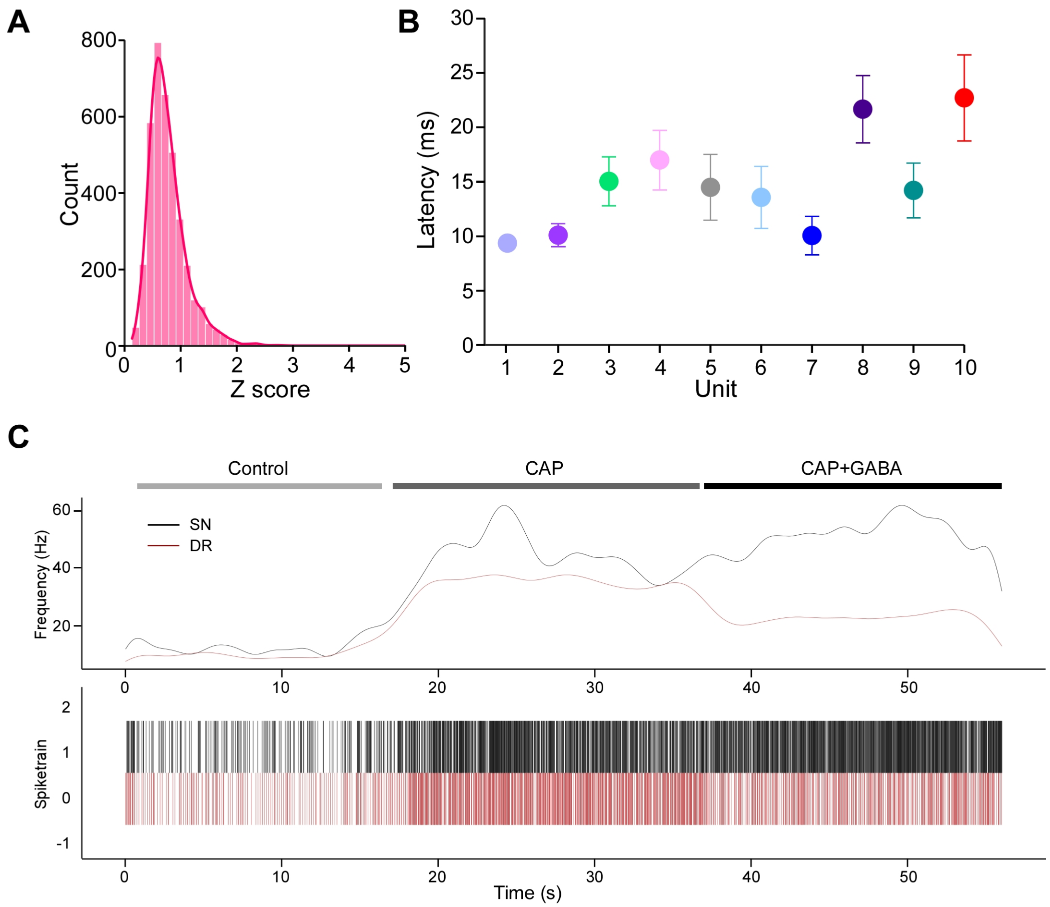


**Supplemental Figure 7**

**Supplemental Figure 7. Additional spike analyses.** (**A**) To rule out contamination of dorsal root units with synchronized firing of another fibre, the mean deviation (represented as a z score) of each waveform in the unit from the mean waveform of the unit was calculated. Any spikes originating from another fibre firing in a temporally correlated way should exhibit a different waveform shape and thus be recognized as an outlier (> 3 z score). The histogram shows a large majority of spikes were within a z score of 3 from the unit means. (**B**) Mean latencies of the DR spikes matched to the units 1 to 10 from the dataset shown in the Suppl. Fig. 6. (**C**) Instantaneous firing frequency of all spike sorted units in the SN and DR.

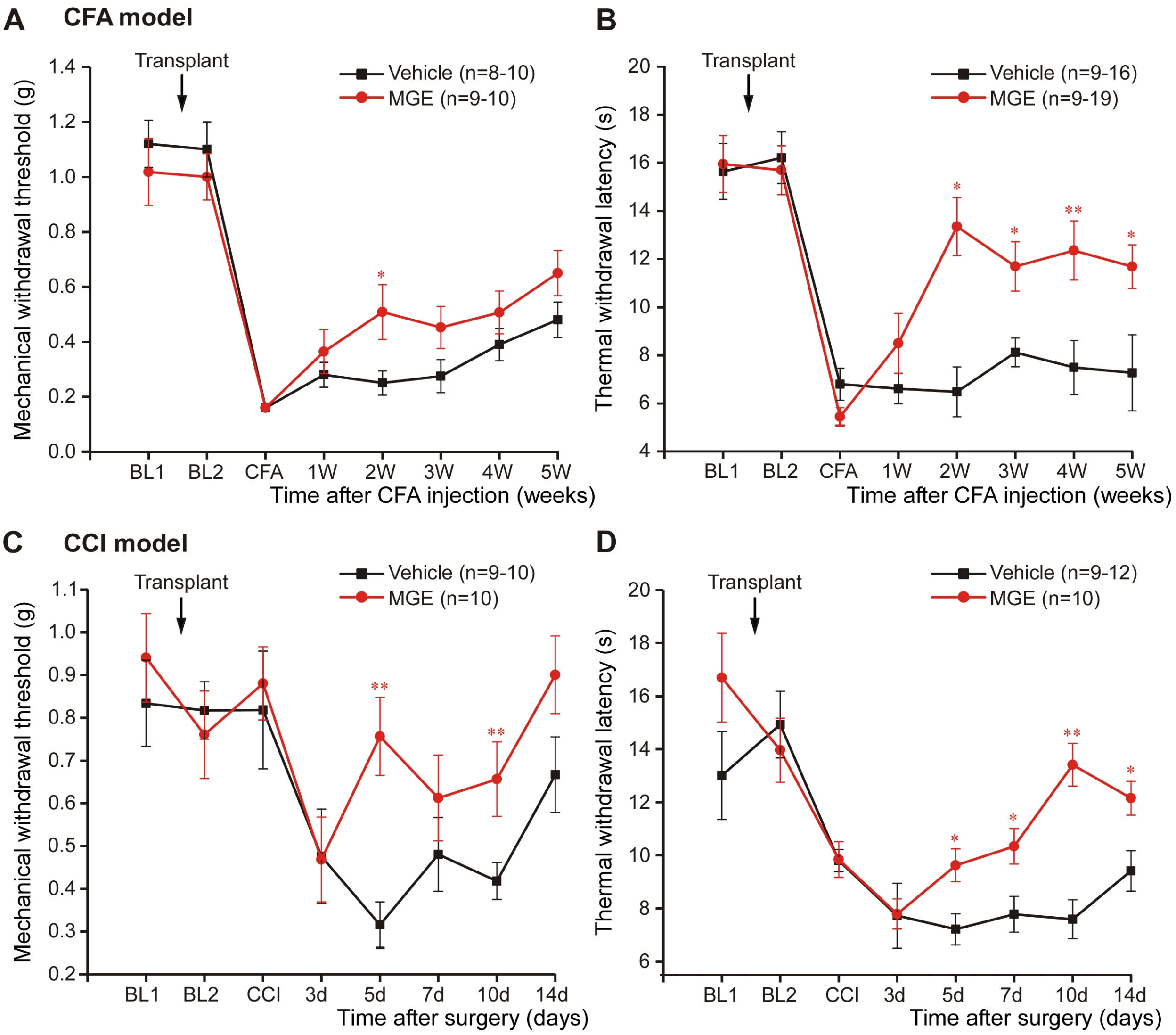


**Supplemental Figure 8**

**Supplemental Figure 8. Transplantation of MGE cells into DRG accelerated the recovery from chronic hyperalgesia.** (**A**, **B**) Mechanical (A) and thermal (B) hyperalgesia caused by hindpaw injection of CFA 2 weeks after MGE cells transplantation into L4 DRG of mice. Mechanical sensitivity was measured using the von Frey method and thermal sensitivity was measured using the Hargreaves method (see Methods). Black symbols denote control mice DRG-injected with saline; red symbols denote MGE-transplanted mice. BL1: baseline before transplantation; BL2: baseline after transplantation; CFA: 1 day after the plantar injection of CFA. Number of experiments (n) is indicated as in each panel (one animal per experiment). (**C**, **D**) experiments similar to A and B, but chronic constriction injury (CCI) neuropathic pain model was performed instead of CFA injection. All labelling is similar to panels A and B. A: Two-factor (MGE vs. vehicle, time after CFA) repeated measures ANOVA: main effects associated with transplantation [$F(1,15)=22.0$; $p<0.01$] and time after CFA [$F(4,12)=2.4$; $p=0.24$]; significant interaction between these factors [$F(4,12)=0.4$; $p=0.83$]. Bonferroni post-hoc test: *significant difference between groups at a given time point ($p<0.05$). B: Two-factor (MGE vs. vehicle, time after CFA) repeated measures ANOVA: main effects associated with transplantation [$F(1,15)=33.2$; $p<0.001$] and time after CFA [$F(4,12)=0.6$; $p=0.69$]; significant interaction between these factors [$F(4,12)=28.2$; $p<0.01$]. Bonferroni post-hoc test: *,**,*** significant difference between groups at a given time point ($p<0.05$, $p<0.01$, $p<0.001$). C: Two-factor (MGE vs. vehicle, time after CCI) repeated measures ANOVA: main effects associated with transplantation [$F(1,17)=11.7$; $p<0.01$] and time after CFA [$F(5,13)=5.6$; $p=0.06$]; significant interaction between these factors [$F(5,13)=1.1$; $p=0.49$]. Bonferroni post-hoc test: *,*** significant difference between groups at a given time point ($p<0.05$, $p<0.001$). B: Two-factor (MGE vs. vehicle, time after CCI) repeated measures ANOVA: main effects associated with transplantation [$F(1,17)=55.1$; $p<0.001$] and time after CFA [$F(5,13)=8.5$; $p<0.05$]; significant interaction between these factors [$F(5,13)=1.5$; $p=0.37$]. Bonferroni post-hoc test: ** significant difference between groups at a given time point ($p<0.01$).

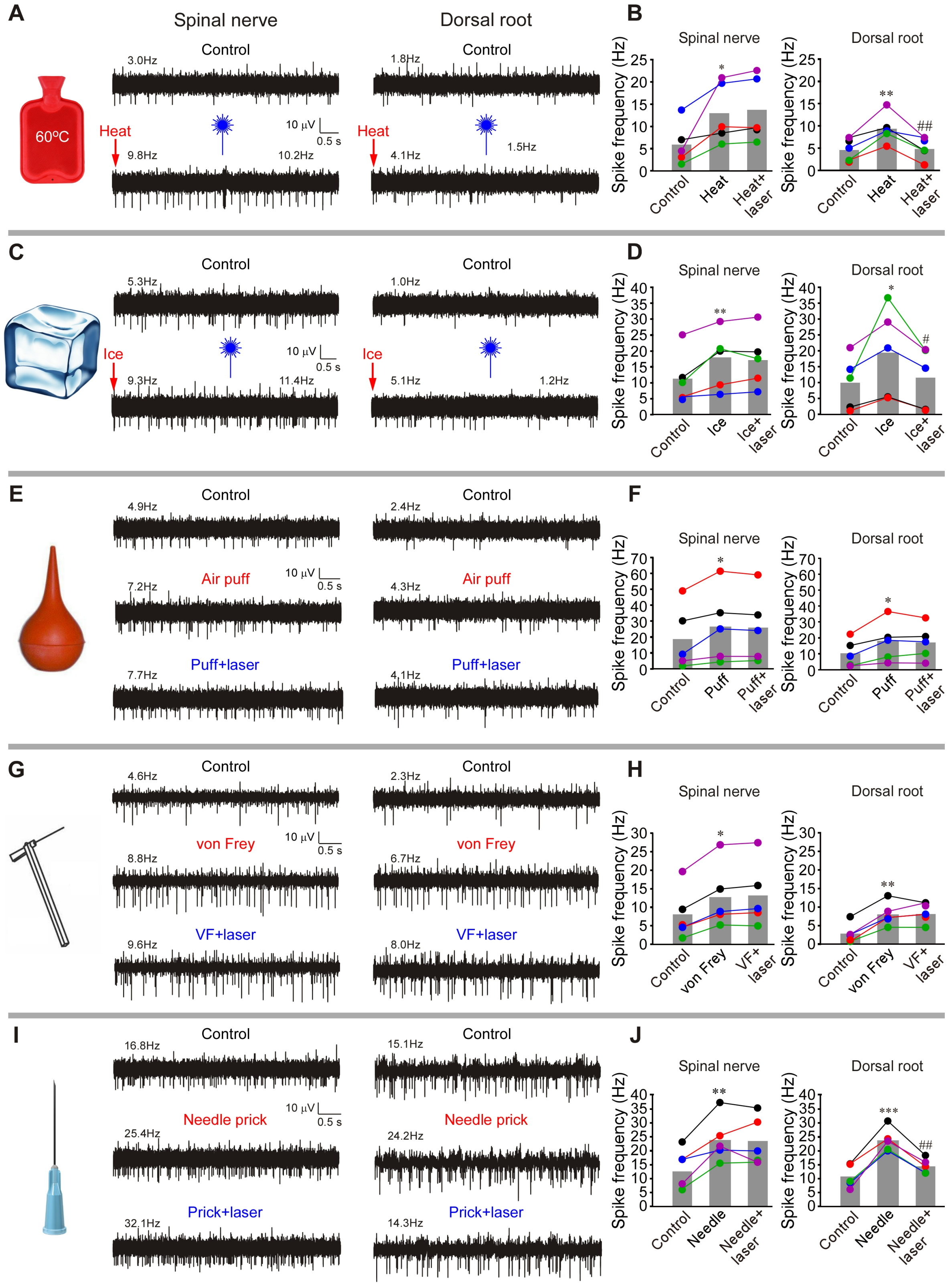


**Supplemental Figure 9**

**Supplemental Figure 9. Optogenetic stimulation of the DRG-transplanted MGE cells enhances filtering of spikes triggered by noxious thermal and mechanical stimulation.** (**A**) Example of *in vivo* recording of the SN and DR activity (similar to these shown in Fig. 1). Stimulation of hindpaw of the MGE-transplanted mice with hot water (60$^{o}$C) increased firing frequency in both SN and DR branches of the nerve (onset of the bottom traces, as compared to basal activity shown in the upper traces). Application of 473 nm laser light to DRG acutely reduced heat-induced firing frequency in DR but not SN (bottom traces). (**B**) Summary for panel A. Two-factor (nerve site, treatment) repeated measures ANOVA: main effect associated with treatment [F(2,7)=10.5; p<0.05]; significant interaction between nerve site and treatment [F(2,7)=12.8; p<0.05]. Bonferroni post-hoc test: *,**significant difference from control (p<0.05, p<0.01); $^{\#\#}$significant difference from heat (p<0.05). (**C**) Similar to A and B but the hindpaw was stimulated with ice cube. (**D**) Summary for panel C. Two-factor (nerve site, treatment) repeated measures ANOVA: significant interaction between nerve site and treatment [F(2,7)=47.8; p<0.01]. Bonferroni post-hoc test: *,**significant difference from control (p<0.05, p<0.01); $^{\#}$significant difference from ice (p<0.01). (**E**) Similar to A and B but the hindpaw was stimulated with air puff. (**F**) Summary for panel E. Two-factor (nerve site, treatment) repeated measures ANOVA: main effect associated with treatment [F(2,7)=11.1; p<0.05]. Bonferroni post-hoc test: *significant difference from control (p<0.05). (**G**) Similar to A and B but the hindpaw was stimulated with sub-threshold von Frey filament (0.4g). (**H**) Summary for panel G. Two-factor (nerve site, treatment) repeated measures ANOVA: main effects associated with treatment [F(2,7)=32.4; p<0.01]. Bonferroni post-hoc test: *,**significant difference from control (p<0.05, p<0.01). (**I**) Similar to A and B but the hindpaw was stimulated with a needle prick. (**J**) Summary for panel I. Two-factor (nerve site, treatment) repeated measures ANOVA: main effect associated with treatment [F(2,7)=235.3; p<0.001]; significant interaction between nerve site and treatment [F(2,7)=38.1; p<0.01]. Bonferroni post-hoc test: *,**significant difference from control (p<0.05, p<0.01); $^{\#\#}$significant difference from needle (p<0.01).

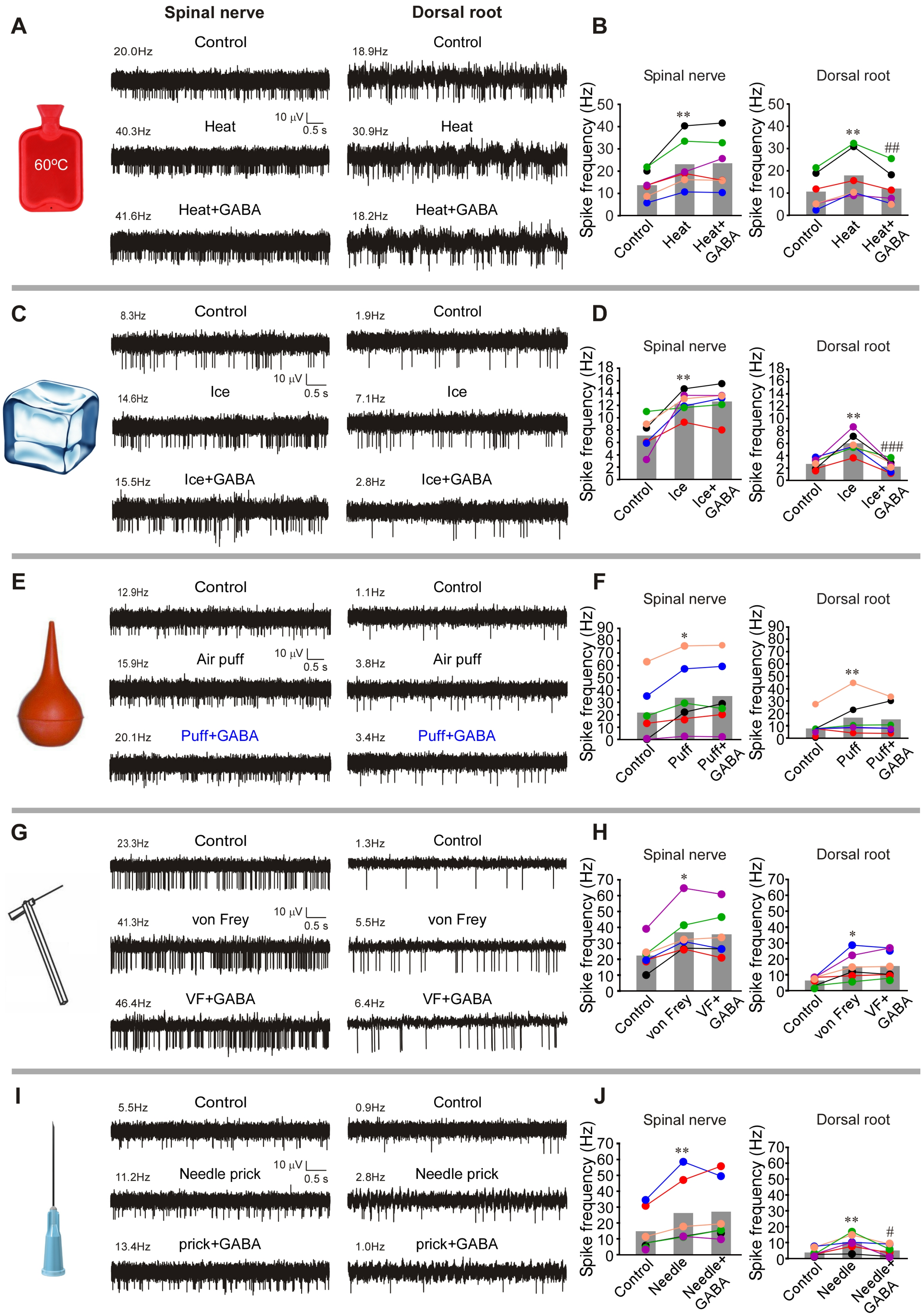


**Supplemental Figure 10**

**Supplemental Figure 10. Activity induced by thermal and mechanical stimulation of the receptive fields is filtered at the DRG.** (**A**) Example of *in vivo* recording of the SN and DR activity (similar to these shown in Fig. 1). Stimulation of hindpaw of the rat with hot water (60oC) increased firing frequency in both SN and DR branches of the nerve (middle traces, as compared to basal activity shown in the upper traces). Application of GABA (200 μM, 3 μl) to DRG reduced heat-induced firing frequency in DR but not SN (bottom traces). (**B**) Summary for panel A. Two-factor (nerve site, treatment) repeated measures ANOVA: main effects associated with nerve site [$F(1,10)=13.4$; $p<0.05$) and treatment [$F(2,9)=8.3$; $p<0.05$]. Bonferroni post-hoc test: **significant difference from control ($p<0.01$); $^{\#\#}$significant difference from heat ($p<0.01$). (**C**) Similar to A and B but the hindpaw was stimulated with ice cube. (**D**) Summary for panel C. Two-factor (nerve site, treatment) repeated measures ANOVA: main effects associated with nerve site [$F(1,10)=150.0$; $p<0.001$) and treatment [$F(2,9)=10.4$; $p<0.05$]; significant interaction between nerve site and treatment [$F(2,9)=13.6$; $p<0.05$]. Bonferroni post-hoc test: **significant difference from control ($p<0.01$); $^{\#\#\#}$significant difference from ice ($p<0.001$). (**E**) Similar to A and B but the hindpaw was stimulated with air puff. (**F**) Summary for panel E. Two-factor (nerve site, treatment) repeated measures ANOVA: main effect associated with treatment [$F(2,9)=7.4$; $p<0.05$]. Bonferroni post-hoc test: $^{*,}$**significant difference from control ($p<0.05$, $p<0.01$). (**G**) Similar to A and B but the hindpaw was stimulated with sub-threshold von Frey filament (4g). (**H**) Summary for panel G. Two-factor (nerve site, treatment) repeated measures ANOVA: main effects associated with nerve site [$F(1,10)=15.0$; $p<0.05$] and treatment [$F(2,9)=11.8$; $p<0.05$]. Bonferroni post-hoc test: *significant difference from control ($p<0.05$). (**I**) Similar to A and B but the hindpaw was stimulated with a needle prick. (**J**) Summary for panel I. Two-factor (nerve site, treatment) repeated measures ANOVA: main effect associated with treatment [$F(2,4)=18.0$; $p<0.01$]; significant interaction between nerve site and treatment [$F(2,4)=7.5$; $p<0.05$]. Bonferroni post-hoc test: **significant difference from control ($p<0.01$); $^{\#}$significant difference from needle ($p<0.05$).

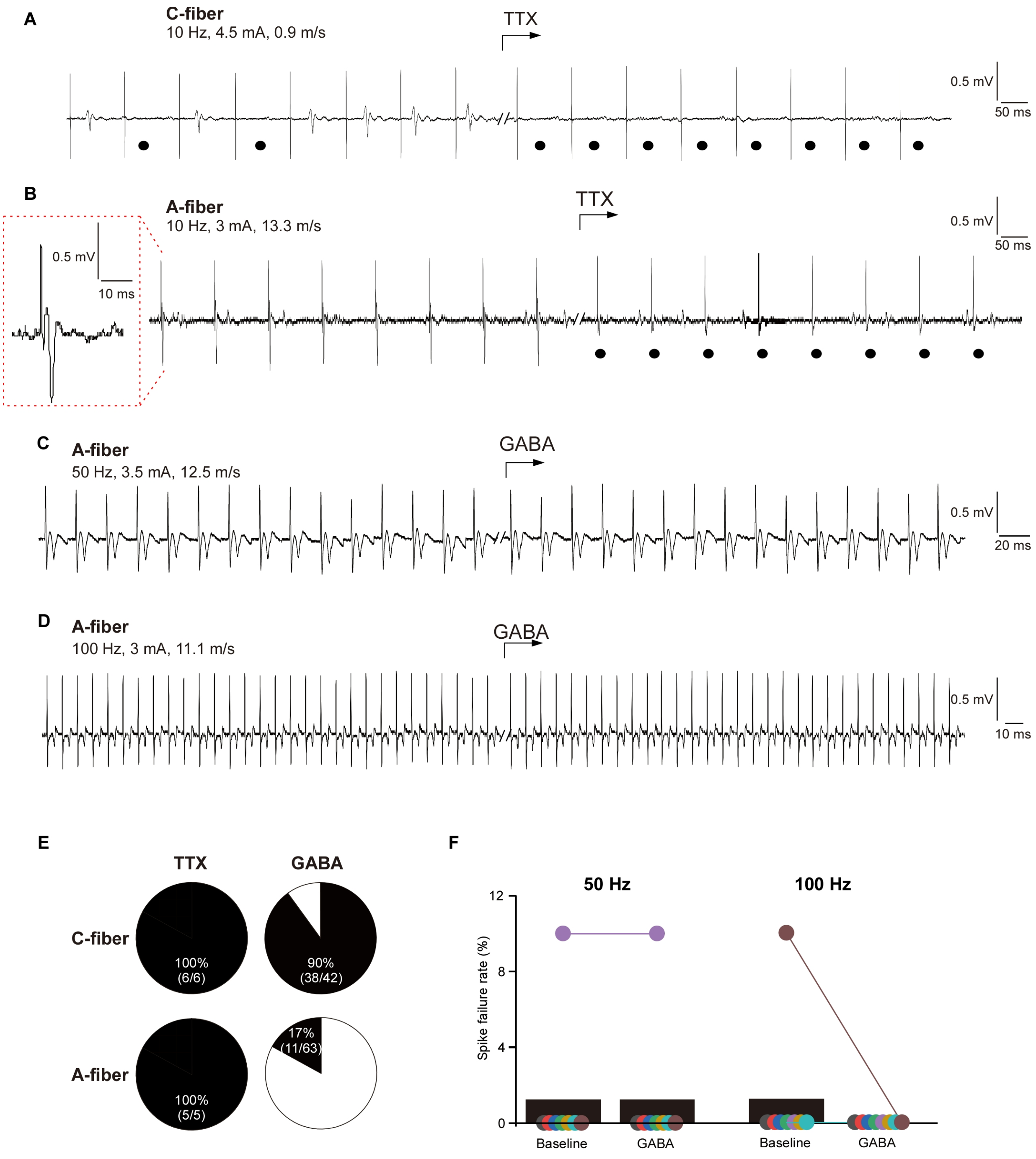


### Supplemental Figure 11

**Supplemental Figure 11. Additional single-unit recordings.** (**A, B**) TTX blocks evoked spikes. Example traces of *in vivo* single unit recording from the DR aspect of a rat C fiber (A) or A fiber (B); stimulus electrode is placed in the spinal nerve. Parameters of stimulation and conduction velocity are indicated above the trace. TTX (1 µM, 3 µl; C) was injected into the DRG using a microsyringe at time points indicated by the bent arrow. Black circles indicate failed spikes. A fiber spike waveform for the A fiber recording is shown on the extended time scale within the dotted red box. (**C, D**). In the A-type fibers spike propagation through the DRG is not affects by GABA even at higher stimulation frequencies: 50 Hz (C) and 100 Hz (D). (**E**) Pie charts summarizing the percentage of C and A fibers in which TTX (from experiments shown in panels A and B) or GABA (data from the experiments shown in main Fig. 5) produced a conduction block. (F) Spike failure rate before and during application of GABA to A fibers when stimulating at 50 and 100 Hz. No significant effects of GABA were found (n=8; Nonparametric Kruskal-Wallis ANOVA).

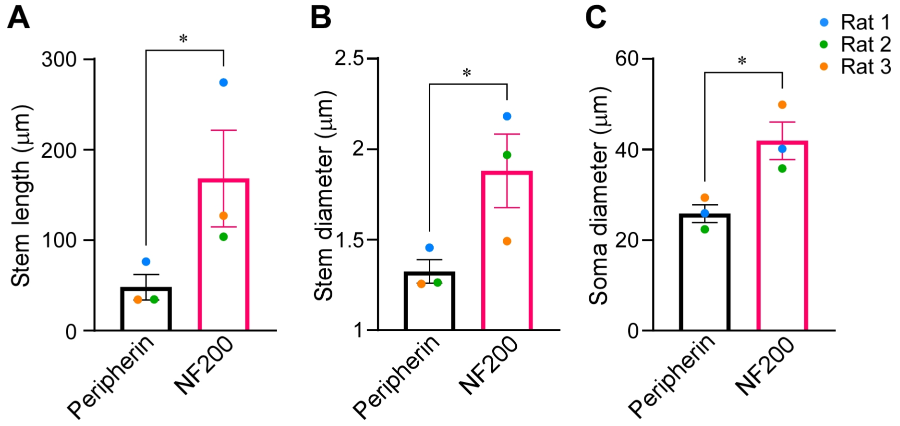


**Supplemental Figure 12**

**Supplemental Figure 12. Additional analysis of initial axon morphology.** Light-sheet microscopy of cleared rat DRG; axon lengths (**A**), stem diameter (**B**) and somatic diameter (**C**) of the peripherin- and NF-200-labelled fibers were analyzed per animal (see also Figure 6C-E). (A) peripherin: 3 animals (rat 1: 5 DRGs, 36 stems; rat 2: 2 DRGs, 9 stems; rat 3: 3 DRGs, 12 stems). NF200: 3 animals (rat 1: 4 DRGs, 22 stems; rat 2: 1 DRG, 3 stems; rat 3: 2 DRGs, 5 stems); paired t-test: $t(2)=3$, $p<0.05$. (B) peripherin: 3 animals (rat 1: 5 DRGs, 36 stems; rat 2: 2 DRGs, 9 stems; rat 3: 3 DRGs, 12 stems). NF200: 3 animals (rat 1: 4 DRGs, 22 stems; rat 2: 1 DRG, 3 stems; rat 3: 2 DRGs, 5 stems); paired t-test: $t(2)=3$, $p<0.05$. (C) peripherin: 3 animals (rat 1: 5 DRGs, 150 somas; rat 2: 2 DRGs, 122 somas; rat 3: 3 DRGs, 155 somas). NF200: 3 animals (rat 1: 4 DRGs, 150 somas; rat 2: 1 DRG, 31 somas; rat 3: 2 DRGs, 75 somas); paired t-test: $t(2)=7$, $p<0.05$.

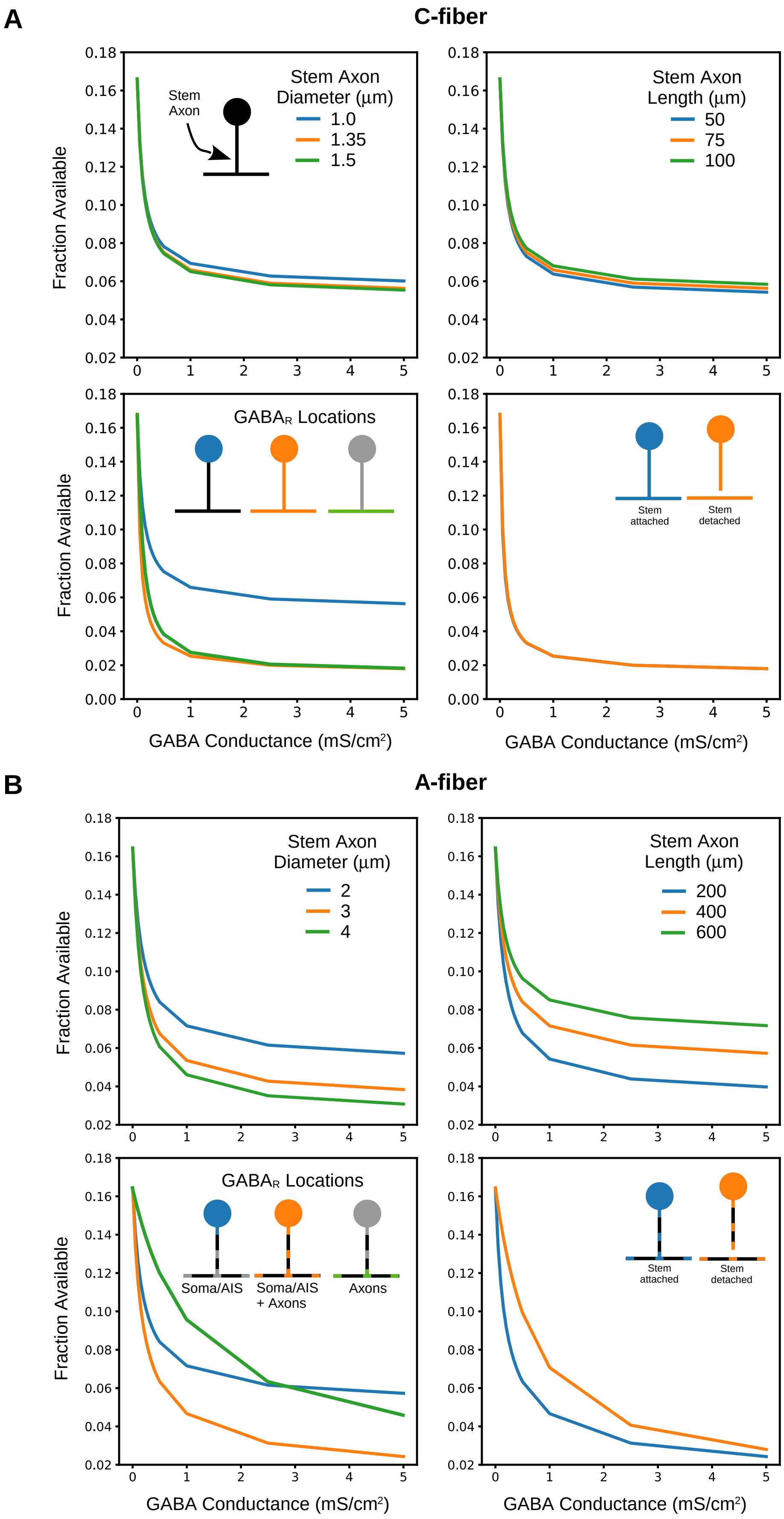


**Supplemental Figure 13**

**Supplemental Figure 13. Parameter space analysis.** (**A**), C-fiber model parameter space analysis. Fraction of available TTX-sensitive Na channels at the T-junction (TJ) in response to steady-state $GABA_A$ receptor activation. For a model with $GABA_A$ receptors restricted to the soma, varying the diameter of the stem axon (length 75 mm; top left panel) or varying stem axon length (diameter 1.35 mm; top right panel) generally had only modest effects on the fraction of available Na channels (i.e. proportion of non-inactivated Na channels). In contrast, expressing $GABA_A$ receptors on the axons, compared with the soma alone, substantially decreased Na channel availability (stem length 75 mm, stem diameter 1.35 mm; bottom left). Connection of the soma and stem to the TJ made no difference when $GABA_A$ receptors were expressed across all compartments, (bottom right). (**B**) A-fiber model parameter space analysis. With $GABA_A$ receptors restricted to the soma, varying stem axon diameter (stem length 4 mm) or diameter (stem diameter 2 mm) substantially affected the available fraction of Na channels (top left and right panels). Likewise, if $GABA_A$ receptors were expressed both in the soma/axon initial segment (AIS) and all nodes of Ranvier, rather than just the soma/AIS alone, fractional availability of Na channels was substantially reduced by $GABA_A$ receptor activation (bottom left). Connection of the soma and stem axon to the TJ provided for a greater effect of $GABA_A$ receptor activation compared to when detached, although this effect was less pronounced with at higher densities of $GABA_A$ receptor activation (bottom right panel).

**Movie S1.** Light sheet microscopy of cleared rat lumbar DRG immunolabeled with NF-200 (red) and peripherin (green).

**Movie S2.** The simple neurite tracer plugin (FIJI) was used to semi-automate stem axons and branch point tracing through a 3D image volume of a DRG. The plugin uses a search algorithm to trace the fluorescently labelled axons between two points. The video shows the example of tracing the stem axon from a peripherin labelled neuron.
